# Increased mutation rate and interlocus gene conversion within human segmental duplications

**DOI:** 10.1101/2022.07.06.498021

**Authors:** Mitchell R. Vollger, William S. DeWitt, Philip C. Dishuck, William T. Harvey, Xavi Guitart, Michael E. Goldberg, Allison N. Rozanski, Julian Lucas, Mobin Asri, The Human Pangenome Reference Consortium, Katherine M. Munson, Alexandra P. Lewis, Kendra Hoekzema, Glennis A. Logsdon, David Porubsky, Benedict Paten, Kelley Harris, PingHsun Hsieh, Evan E. Eichler

## Abstract

Single-nucleotide variants (SNVs) within segmental duplications (SDs) have not been systematically assessed because of the difficulty in mapping short-read sequence data to virtually identical repetitive sequences. Using 102 phased human haplotypes, we constructed 1:1 unambiguous alignments spanning high-identity SDs and compared the pattern of SNVs between unique and SD regions. We find that human SNVs are elevated 60% in SDs compared to unique regions. We estimate that at least 23% of this increase is due to interlocus gene conversion (IGC) with >7 Mbp of SD sequence converted on average per human haplotype. We develop a genome-wide map of IGC donors and acceptors, including 498 acceptor and 454 donor hotspots affecting the exons of ~800 protein-coding genes. The latter includes 171 genes that have “relocated” on average 1.61 Mbp in a subset of human haplotypes. Using a coalescent framework, we show that SD regions are evolutionarily older when compared to unique sequences with most of this signal originating from putative IGC loci. SNVs within SDs, however, also exhibit a distinct mutational spectrum where there is a 27.1% increase in transversions that convert cytosine to guanine or the reverse across all triplet contexts. In addition, we observe a 7.6% reduction in the frequency of CpG associated mutations when compared to unique DNA. We hypothesize that these distinct mutational properties help to maintain an overall higher GC content of SD DNA when compared to unique DNA, and we show that these GC-favoring mutational events are likely driven by GC-biased conversion between paralogous sequences.

## INTRODUCTION

The landscape of human single-nucleotide variants (SNVs) has been well understood for more than a decade in large part due to large-scale efforts such as the HapMap and 1000 Genomes Project (McCarroll et al. 2006; 1000 Genomes Project Consortium et al. 2015, 2012; Sudmant et al. 2015; 1000 Genomes Project Consortium et al. 2010). While these consortia helped establish the genome-wide pattern of SNVs (as low as 0.1% in frequency) and linkage disequilibrium based on sequencing and genotyping thousands of human genomes, not all parts of the human genome could be equally ascertained. Approximately 10-15% of the human genome (1000 Genomes Project Consortium et al. 2012) has remained inaccessible to these types of analyses either because of gaps in the human genome or, more frequently, the low mapping quality associated with aligning short-read whole-genome sequencing data. This is because short-read sequence data are of insufficient length (<300 bp) to unambiguously assign reads and, therefore, variants to specific loci (Sudmant et al. 2010). While certain classes of large highly identical repeats (e.g., alpha-satellites in centromeres) were readily recognized, others, especially segmental duplications (SDs) (J. A. Bailey et al. 2001; Jeffrey A. Bailey et al. 2002) and their associated 859 genes (Vollger et al. 2022), in euchromatin were much more problematic. This led to the misclassification of paralogous sequence variants (PSVs) as SNVs (Hurles 2002) and, as a result, high-identity SDs became blacklisted from subsequent genomic analyses (Zook et al. 2019; 1000 Genomes Project Consortium et al. 2015; Amemiya, Kundaje, and Boyle 2019). This exclusion has translated into a fundamental lack of understanding in mutational processes precisely in regions predicted to be more mutable due to the action of ectopic or interlocus gene conversion (IGC) (Teshima and Innan 2012). Leveraging high-quality phased genome assemblies generated as part of the Human Pangenome Reference Consortium (HPRC)(Liao et al. 2022), we compare the SNV landscape of duplicated and unique DNA in the human genome.

## RESULTS

### Strategy and quality control

Unlike previous SNV discovery efforts, which cataloged SNVs based on the alignment of sequence reads, our strategy was assembly driven. We focused on the comparison of 102 haplotype-resolved genomes generated as part of the HPRC (n=94) or other efforts (n=8) (Ebert et al. 2021; Ebler et al. 2022; IHGSC 2001; Schneider et al. 2017; Nurk et al. 2022; Liao et al. 2022) where phased genome assemblies had been assembled using high-fidelity (HiFi) long-read sequencing (Cheng et al. 2021; Jarvis et al. 2022).The extraordinary assembly contiguity of these haplotypes (contig N50>40 Mbp) provided an unprecedented opportunity to align large swathes (>1 Mbp) of the genome, including large SD repeats anchored by megabases of synteny (see below). Notwithstanding, SDs are frequently a source of misassembly even among phased genome assemblies (Ebert et al. 2021), so we performed a number of additional validation experiments to assess the quality of HPRC SDs and quantify SD missassembly errors. Using the telomere-to-telomere (T2T) reference (v1.1) as a guide of completeness and the HPRC callset of potentially unreliable regions (Liao et al. 2022), we determined that, on average, only 1.64 Mbp (1.37%) of the analyzed SD sequence was suspect due to abnormal read coverage (Supplement). A comparison of copy number variable SD regions in the haplotype-resolved assembly with orthogonal short-read estimates showed a high degree of copy number correlation (Pearson R^2 = 0.97, Fig. S1). Only three large (>140 kbp) and highly identical (>99.3%) SDs were consistently discordant by short-read copy number variant estimates (Fig. S2). Similarly, we leveraged orthogonal ONT data from the same samples and applied a recently developed method that assesses read depth based on mapping of distance between unique *k*-mers (Dishuck et al. 2022). This haplotype-specific analysis confirmed the copy number and integrity of >94% of all tested SDs. As a final control for potential haplotype-phasing errors, we generated deep ONT and HiFi data from a second hydatidiform mole (CHM1) where a single paternal haplotype was present (Chaisson et al. 2015; Vollger et al. 2020) and show that across our many analyses the results from the CHM1 Verkko assembly are consistent with individual haplotypes obtained from diploid HPRC samples produced by trio-hifiasm (Cheng et al. 2021; Rautiainen et al. 2022) (Figs. S3-S4). We conclude that the vast majority (>95%) of SDs analyzed were accurately assembled from multiple human genomes allowing the pattern of SNV diversity to be systematically interrogated.

### Increased single-nucleotide variation in SD regions

To assess SNVs, we limited our analysis to portions of the genome where a one-to-one orthologous relationship could be unambiguously assigned (as opposed to regions with extensive copy number variation). Using the T2T-CHM13 reference genome, we aligned the HPRC haplotypes requiring alignments to be a minimum of 1 Mbp in length and carry no structural variation events greater than 10 kbp (Supplemental Methods). While the proportion of haplotypes compared for any locus varied (Fig. 1a), the procedure allowed us to establish, on average, 120.2 Mbp 1:1 fully aligned sequence per genome for SD regions out of a total of 217 Mbp from the finished human genome (T2T-CHM13 v1.1). We repeated the analysis for “unique” regions of the genome (single-copy regions of the genome) and recovered by comparison 2,508 Mbp as 1:1 alignments (Fig. 1a). All downstream analysis was then performed using this orthologous alignment set. We first compared the SNV diversity between unique and duplicated regions excluding suboptimal alignments mapping to tandem repeats or homopolymer stretches. Overall, we observe a significant 60% increase in SNVs in SD regions (Supplemental Methods, Pearson’s Chi-squared test with Yates’ continuity correction p<2.2e-16, Fig. 1b,d). Specifically, we observe an average of 15.3 SNV per 10 kbp versus 9.57 SNVs per 10 kbp for unique sequences (Fig. 1d). An empirical cumulative distribution (eCDF) comparing the divergence of individual 10 kbp windows between SD and unique sequence confirms that this is a general property and not driven simply by outliers. The eCDF shows that more than half of the SD sequences are more divergent than their unique counterparts (Fig. 1b).

**Figure 1.**
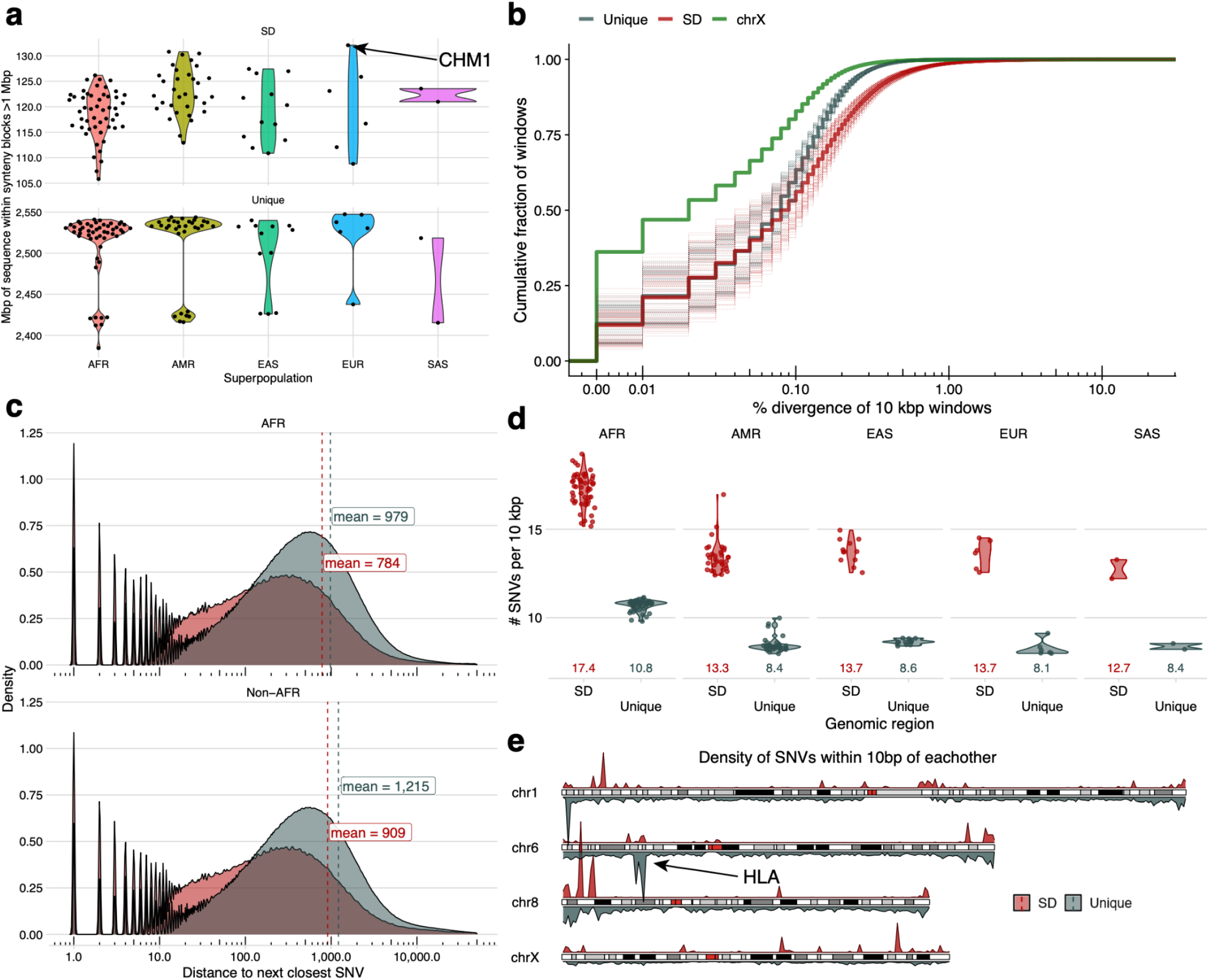
Increased single-nucleotide variation in SDs. **a)** The portion of the haplotype-resolved human genome aligned in Mbp for SD regions (top) and unique regions (bottom) of the genome based on 1:1 syntenic blocks per human superpopulation (color). **b)** Empirical cumulative distribution showing the average % sequence divergence of 10 kbp windows within the syntenic regions stratified by unique (gray), SD (red), and the X chromosome (green). Dashed lines represent individual haplotypes and thick lines represent the average trend of all the data. **c)** Distribution of the average distance to the next closest SNV in SD (red) and unique (gray) space separating African (top) and non-African (bottom) samples. Dashed vertical lines are drawn at the mean of each distribution. **d)** Average number of SNV per 10 kbp window in SD (red) vs. unique (gray) space by superpopulation and with mean value displayed underneath each violin. **e)** Density of SNVs within 10 bp of each other for SD space (top, red) and unique regions (bottom, gray) for chromosomes 1, 6, 8, and X showing the relative density of known (e.g., HLA) and new hotspots of single-nucleotide variation.

Previous publications have shown that African haplotypes are genetically more diverse having on average ~20% more variant sites against the reference per sample compared to non-Africans (1000 Genomes Project Consortium et al. 2015). To confirm this observation in our data we examined the number of SNV per 10 kbp of unique sequence in Africans versus non-Africans and (Fig. 1c,d) observed a 27% (10.8/8.5) excess in Africans. As a result, among Africans we see that the average distance between SNVs (979 bp) is 19.4% closer than in non-Africans (1,215 bp) as expected (1000 Genomes Project Consortium et al. 2015; Sudmant et al. 2015; IHGSC 2001). African genomes also show increased variation within SDs but it is less pronounced with an average distance of 784 bases between consecutive SNVs as compared to 909 bases in non-Africans. Although elevated in Africans, SNV density is higher in SD sequence across populations and these properties are not driven by a few sites but, once again, are a genome-wide feature. We put forward three possible hypotheses to account for this increase although note these are not mutually exclusive: 1) SDs have unique mutational mechanisms that increase SNVs, 2) SDs have a deeper average coalescence than unique parts of the genome, and 3) differences in sequence composition (i.e., GC richness) make SDs more prone to particular classes of mutation.

### Signatures of interlocus gene conversion (IGC)

A possible explanation for increased diversity in SDs is IGC where sequence that is orthologous by position no longer shares an evolutionary history because a paralog from a different location has “donated” its sequence via ectopic template-driven conversion (Bosch et al. 2004). To identify regions of IGC, we developed a method that compares two independent alignment strategies to pinpoint regions where the orthologous alignment of an SD sequence is inferior to an independent alignment of the sequence without flanking information (Fig. S5, Supplemental Methods). The orthologous alignment we refer to as an alignment by position (ABP) while the second we refer to as an alignment by sequence (ABS)—as it is the best sequence alignment but is no longer necessarily at the orthologous location. ABP was determined using the previously established >1 Mbp windows of 1:1 alignment and ABS was determined by extracting 1 kbp windows (with a 100 bp slide) of SD sequence and allowing them to align without considering flanking sequence information. We defined candidate IGC as positions where the ABS alignment was distinct and improved compared to the ABP alignment, with the acceptor site defined by the ABP alignment and the donor site defined by the ABS alignment. Furthermore, we show that our high-confidence IGC calls (20+ supporting SNVs) have strong overlap with other methods for identifying IGC in regions known to have IGC (Supplemental Information) (Sawyer 1989, n.d.; Hsieh et al. 2020). Using this approach, we created a genome-wide map of putative large IGC events for all the HPRC haplotypes where 1:1 orthologous relationships could be established (Fig. 2).

**Figure 2.**
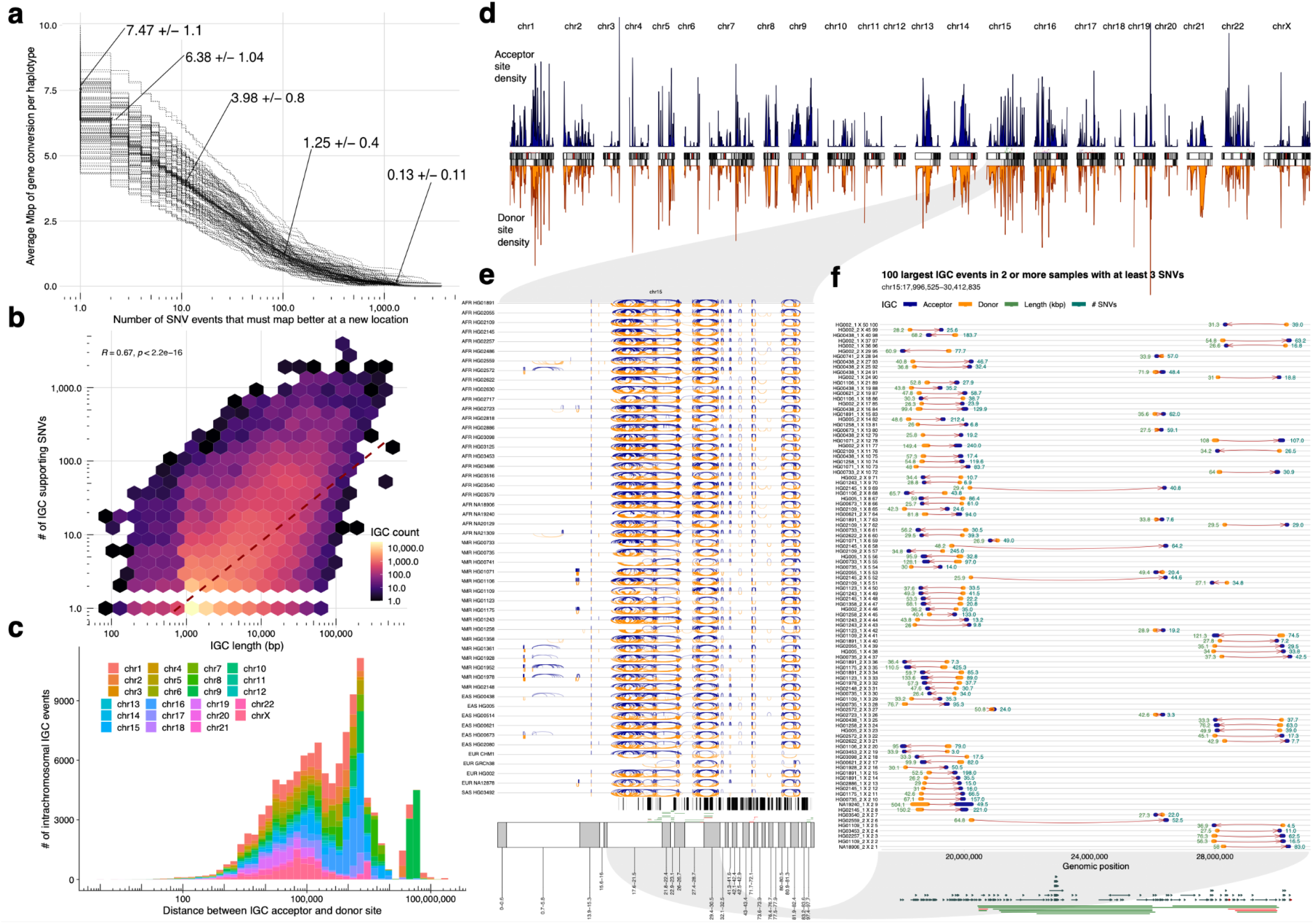
Candidate interlocus gene conversion events. **a)** Mbp of interlocus gene conversion (IGC) observed in HPRC haplotypes, as a function of the minimum number of SNVs that support the IGC call. Dashed lines represent individual haplotypes and the solid line represents the average. **b)** Correlation between IGC length and the number of supporting SNVs. **c)** Distribution of the distance between IGC acceptor and donor sites for intrachromosomal events by chromosome. **d)** Density of IGC acceptor (top, blue) and donor (bottom, orange) sites across the “SD genome”. The SD genome consists of all major SD regions (>50 kbp) minus the intervening unique sequences. **e)** All intrachromosomal IGC events on the human haplotypes analyzed for chromosome 15. Arcs drawn in blue (top) have the acceptor site on the left-hand side and the donor site on the right. Arcs drawn in orange (bottom) are arranged oppositely. Protein-coding genes are drawn as vertical black lines above the ideogram, and large duplication (blue) and deletion (red) events associated with human diseases are drawn as horizontal lines just above the ideogram. **f)** Zoom of the 100 largest IGC events on chromosome 15 between 17 and 31 Mbp.

Across all 102 haplotypes, we observe 121,631 putative IGC events for an average of 1,193 events per human haplotype (Table S1). Of these events, 17,949 are rare and restricted to a single haplotype (singletons) while the remaining events are observed in multiple human haplotypes grouping into 14,663 distinct events (50% reciprocal overlap at both the donor and acceptor site). In total, we estimate there is evidence for 32,612 different IGC events (Table S2) at least among the SD regions that are currently assessed. Considering the redundant IGC callset (n=121,631), the average length of IGC observed in our data is 6.26 kbp with the largest event we observe being 504 kbp (Fig. S6). On average, each IGC event has 13.3 SNVs that support the conversion event and 2.03 supporting SNVs per kbp, and as expected there is strong correlation (R=0.63, pearsons) between the length of the events and supporting SNVs. We further stratify these results by callset, minimum number of supporting SNVs, and haplotype (Table S3).

On average, we identify 7.47 Mbp of sequence per haplotype affected by IGC. Overall, 33.8% (60.77 Mbp / 180.0 Mbp) of the analyzed SD sequence is affected by IGC in at least one human haplotype. Furthermore, among all SDs covered by at least 20 assembled haplotypes, we identify 498 acceptor and 454 donor IGC hotspots with at least 20 distinct IGC events (Table S4, Fig. 2d). IGC hotspots are more likely to associate with higher copy number SDs compared to a random sample of SD windows of equal size (median of 9 overlaps compared to 3, one-sided Wilcoxon rank sum test p<2.2*e-16). Therefore, IGC appears to be preferentially located in higher copy number repeats. The number of distinct IGC events at a particular locus is moderately correlated with the log of the copy number over the same window (Pearson’s R = 0.2, p<2.2e-16). IGC hotspots also preferentially overlap higher identity duplications (median 99.4%) compared to randomly sampled windows (median 98.0%, one-sided Wilcoxon rank sum test p<2.2*e-16).

These events intersect 1,179 protein-coding genes and of these genes 799 have at least one coding exon affected by IGC (Table S5). As a measure of functional constraint, we used the probability of being loss-of-function intolerant (pLI) for each of the 799 genes (Lek et al. 2016). Among these, 314 have no pLI score likely due to the limitations of mapping short-read data from population samples. Of the remaining genes we identify 38 with a pLI greater than 0.5, including genes important in disease (*F8*, *HBG1*, *C4B*) and evolution (*NOTCH2*, *TCAF*). Of the genes with high pLI scores, 12 are the acceptor site for at least 50 IGC events including *CB4*, *NOTCH2*, and *OPNL1W*. We identify a subset of 418 nonredundant IGC events that are predicted to move the entirety of a gene body to a “new location” in the genome. As a result 171 different protein-coding genes with at least 2 exons and 200 coding base pairs are repositioned by IGC in a subset of human haplotypes (Table S6). These gene-repositioning events are large (average 26 kbp; median 16.7 kbp) and supported by a high number of SNVs (average 64.7; median 15.3 SNVs) suggesting they are unlikely mapping artifacts. Remarkably, these IGC gene relocations move the reference gene model on average 1.66 Mbp (median 216 kbp) from its original location with potential differential consequences with respect to regulation. These include several disease-associated genes (e.g., *TAOK2*, *C4A*, *C4B*, *PDPK1*, *IL27*) and genes that have alluded complete characterization due to their duplicative nature (Richter et al. 2019; Sekar et al. 2016; Shahi et al. 2020; Pietri et al. 2013).

### Evolutionary age of SDs

Notwithstanding the potential of IGC to reshape the organization of the genome for specific human haplotypes, we estimate that IGC contributes modestly to the significant increase of human SNV diversity in SDs. For example, if we apply the least conservative definition of IGC (1 supporting SNV) and exclude all putative IGC events from the human haplotypes, we estimate it accounts for only 23% of the increase in single-nucleotide variation within SDs (Fig. S7), though IGC from donor haplotypes not observed in our dataset could potentially increase this estimate. An alternative explanation may be that the SDs are evolutionarily older, perhaps due to reduced selective constraint on duplicated copies (Force et al. 1999; Conant and Wagner 2003). To test if SD sequences appear to have a deeper average coalescence than unique regions, we constructed a high-quality locally phased assembly (hifiasm v0.15.2) of a chimpanzee genome (*Pan troglodytes*) to calibrate age since the time of divergence and to distinguish ancestral versus derived alleles in human SD regions (Supplemental Methods). Constraining our analysis to syntenic regions between human and chimpanzee (Supplemental Methods), we characterized 4,316 SD regions (10 kbp in size) where we had variant calls from at least 50 human and one chimpanzee haplotype. We selected at random 9,247 analogous windows from unique regions for comparison. We constructed a multiple sequence alignment for each window and estimated the time to the most recent common ancestor (TMRCA) for each 10 kbp window independently. We infer that SDs are significantly older than the corresponding unique regions of similar size (Fig. S8) (one-sided Wilcoxon rank sum test p value=4.3e-14), assuming that mutation rates have remained constant over time within these regions since the human/chimpanzee divergence. The TMRCAs inferred from SD regions are on average 22% more ancient when compared to unique regions (650 vs. 530 thousand years ago (kya)) but only a 5% difference is noted when comparing the median (520 vs. 490 kya). However, this effect all but disappears (only a 0.2% increase) after excluding windows classified as IGC (Figs. S9-S10, one-sided Wilcoxon rank sum test p = 0.05; mean TMRCA_unique_ = 0.528, mean TMRCA_SD_ = 0.581, median TMRCA_unique_= 0.495, median TMRCA_SD_= 0.496).

### SD mutational spectra

As a third possibility, we considered potential differences in the sequence context of unique and duplicated DNA. It has been recognized for almost two decades that human SDs are particularly biased toward Alu repeats and GC-rich DNA of the human genome (Jeffrey A. Bailey, Liu, and Eichler 2003; Zhang et al. 2005; Nakken et al. 2009). Notably, among the SNVs within SDs we observed a significant excess of transversions (1.78 Ti/Tv) when compared to unique sequence (2.06 Ti/Tv) ((p<2.2e-16, Pearson’s Chi-squared test with Yates’ continuity correction). Increased mutability of GC-rich DNA is expected and may explain, in part, the increased variation in SDs and transversion bias (Kiktev et al. 2018; Zhu et al. 2014; Duncan and Miller 1980). Using a more complete genome, we compared the GC composition of unique and duplicated DNA specifically for the regions considered in this analysis. We find that on average that 42.4% of the analyzed SD regions are guanine or cytosine (43.0% across all SDs) when compared to 40.8% of the unique portions of the genome (p-value < 2.2e-16, one-sided t-test). Interestingly, this enrichment drops slightly (41.8%) if we exclude IGC regions. Consequently, we observe an increase of all GC-containing triplets in SD sequences compared to unique regions of the genome (Fig. 4a). Furthermore, the enrichment levels of particular triplet contexts in SD sequence correlate with the mutability of the same triplet sequence in unique regions of the genome (Pearson R = 0.77, p = 2.4e-07, Fig. 4b). This effect is primarily driven by CpG-containing triplets, which are enriched between 14 and 30% in SD sequences; however, there is still a (weaker) correlation in the non-CpG-containing triplets (Pearson R = 0.2). Extrapolating from the mutational frequencies seen in unique sequences, we estimate that there is 3.21% more variation with SDs due to their sequence composition alone.

**Figure 3.**
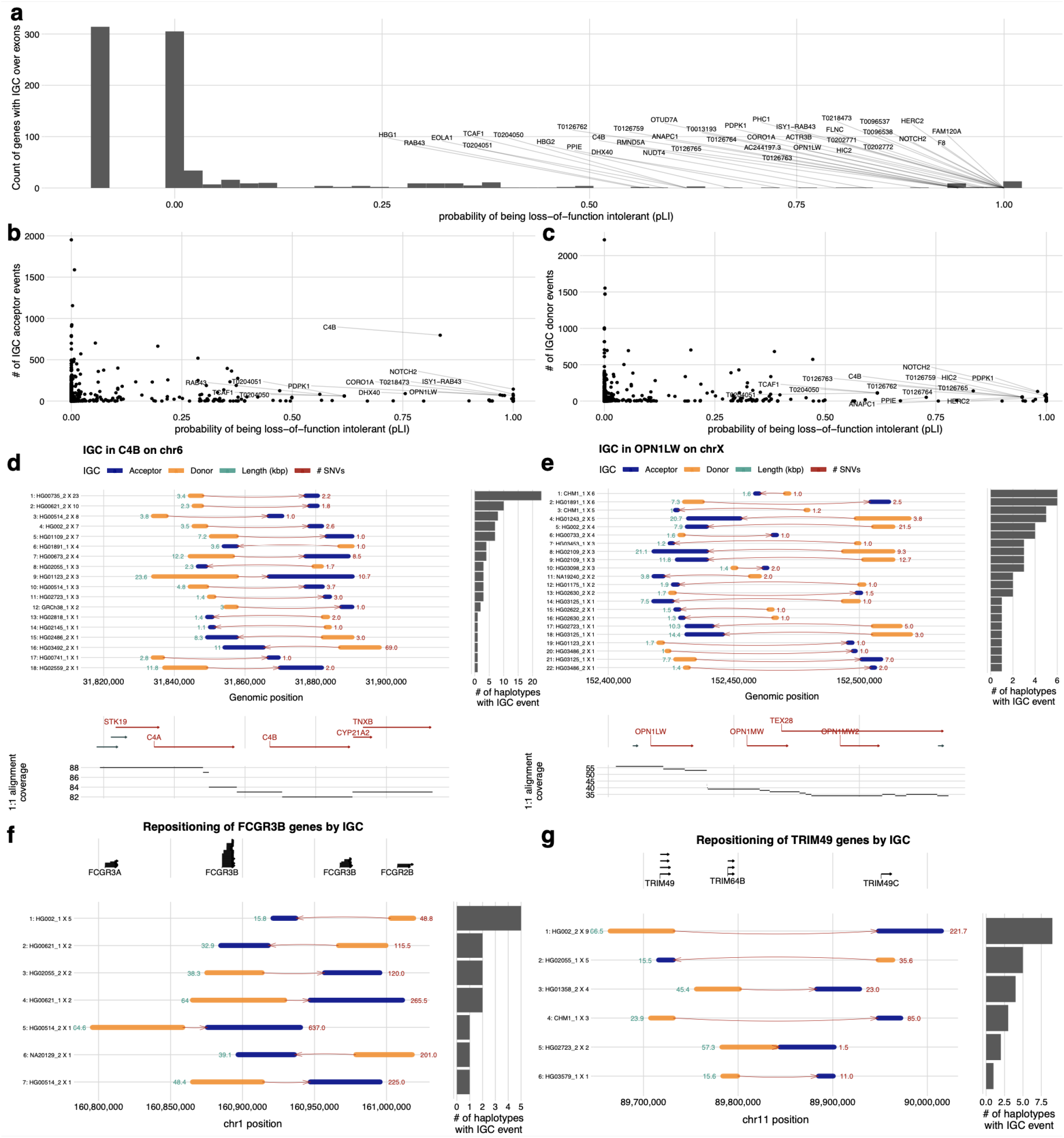
Protein-coding genes affected by IGC. **a)** Number of IGC events intersecting exons of protein-coding genes as a function of a gene’s probability of loss of function (pLI). **b,c)** Number of times a gene exon acts as an acceptor (**b**) or a donor (**c**) of an IGC event. **d)** IGC events at the complement factor locus, *C4A* and *C4B* and **e)** the opsin middle and long wave-length sensitive genes (*OPN1MW* and *OPN1LW* locus). Predicted donor (orange) and acceptor (blue) segments by length (green) and average number of supporting SNVs (red) are shown. The number of human haplotypes supporting each configuration are depicted by the histograms to the right. **f,g)** IGC events that reposition entire gene models for the *FCGR* (**f**) and the *TRIM* (**g**) loci. For additional examples of gene-associated IGC, see Supplementary Figures.

**Figure 4.**
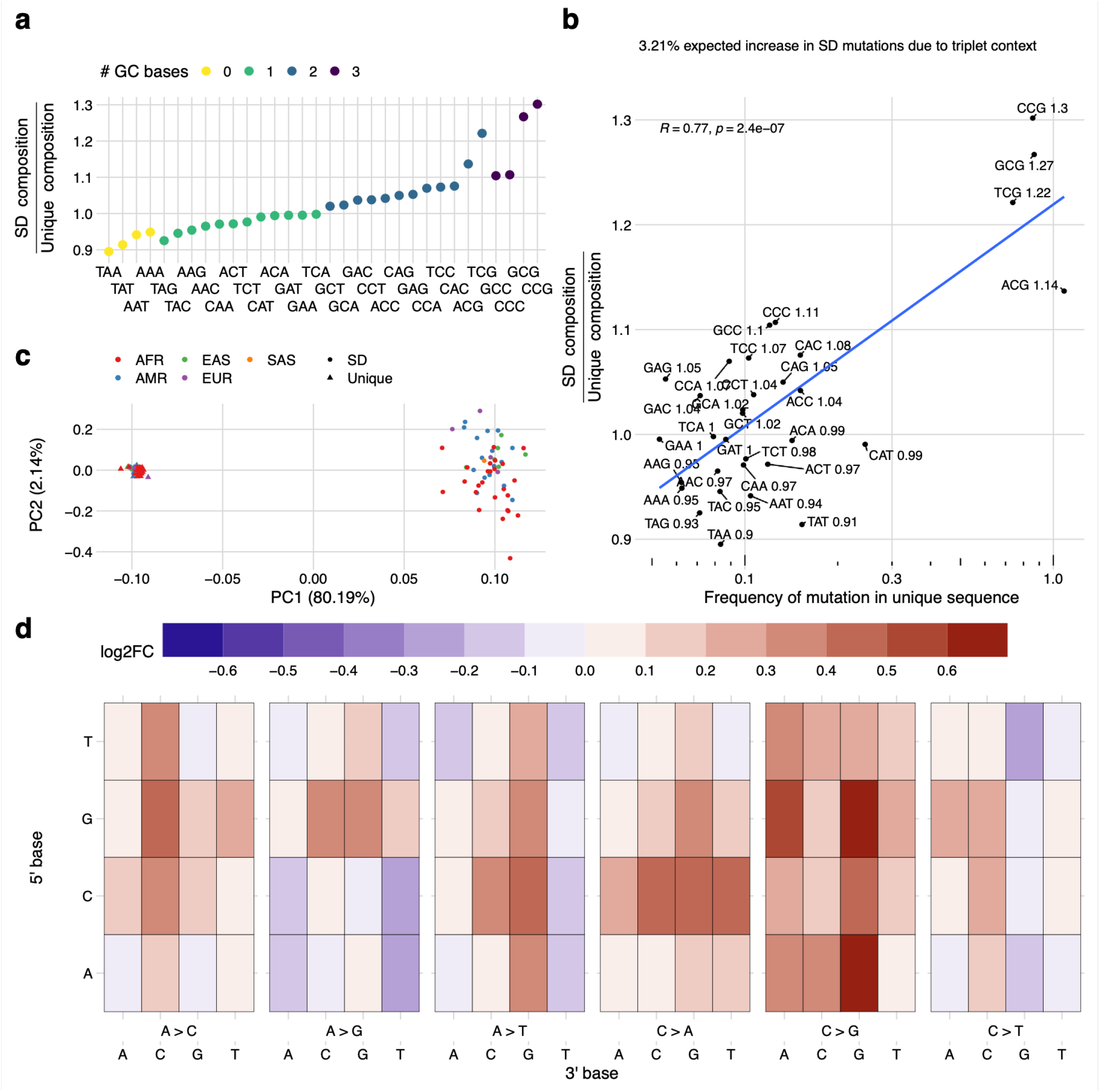
Sequence composition and mutational spectra of SD SNVs. **a)** Compositional increase in GC-containing triplets (3mers) in SDs versus unique regions of the genome (colored by GC content). **b)** Shows a correlation between the enrichment of certain triplets in SDs compared to the mutability of that triplet in unique regions of the genome. Mutability is defined as the sum of all SNVs that change a triplet divided by the total count of that triplet in the genome. The enrichment ratio of SD over unique is indicated in text next to each triplet sequence. **c)** PCA of the mutational spectra of triplets in SD (circles) vs. unique (triangles) regions polarized against a chimpanzee genome assembly and colored by the continental superpopulation of the sample. **d)** Heatmap of the log-fold change in the triplet-normalized mutation frequency between SDs and unique sequences for the same mutational event.

To further investigate the changes in GC content and their effect on variation in SDs, we compared the triplet mutational spectra of SNVs from unique and duplicated regions of the genome to determine if there were significant shifts in the predominant modes of mutation (DeWitt 2020; Carlson, DeWitt, and Harris 2020). We considered all possible triplet changes (3-mers), first quantifying the number of ancestral GC bases and triplets in SDs (Fig. 4a) (Jiang et al. 2007; Harris 2015; DeWitt 2020). A PCA of these normalized mutational spectra shows clear discrimination (Fig. 4c) for both unique and SD regions (PC1) beyond that of African and non-African diversity. We observe several differences when comparing the triplet-normalized mutation frequency of particular mutational events in SD and unique sequences (Fig. 4d). Most notable is a 7.6% reduction in CpG transition mutations–the most predominant mode of mutation in unique regions of the genome due to spontaneous deamination of methylated CpGs (Duncan and Miller 1980) (Table S7).

The most striking changes in mutational spectra in SD sequences are: a 27.1% increase in C>G mutations, a 15.3% increase in C>A mutations, and a 10.5% increase in A>C mutations. C>G mutations are associated with double-stranded breaks in humans and some other apes (Jónsson et al. 2017; Rahbari et al. 2016; Goldmann et al. 2018; Gao et al. 2019). This effect becomes more pronounced (+40.4%) in our candidate IGC regions consistent with previous observations showing increases in C>G mutations in regions of non-crossover gene conversion and double-strand breaks (Elliott et al. 1998; Jónsson et al. 2017; Gao et al. 2019). However, the increase remains in SD regions without IGC (+20.0%) perhaps due to extensive non-allelic homologous recombination (NAHR) associated with SDs or undetected IGC events (Sudmant et al. 2015; Chaisson et al. 2015; Huddleston et al. 2017; Chaisson et al. 2019; Ebert et al. 2021).

To further investigate the potential effect of GC-biased gene conversion (gBGC) on the mutational spectra within SDs, we measured the frequency of (A,T) > (G,C) mutations in SD regions with evidence of IGC to see if cytosine and guanine bases are being preferentially maintained as might be expected in regions undergoing gBGC. If we measure the frequency of (A,T) > (C,G) in windows where at least one haplotype has evidence of IGC, then we see the frequency is 4.7% higher than in unique regions of the genome; interestingly, in SDs without IGC this rate is reduced compared to unique sequence (−3.5%). Additionally, there is a 5.8% reduction in (G,C) > (A,T) bases consistent with IGC preferentially restoring CG bases that have mutated to AT bases via gBGC. These results indicate that gBGC between paralogous sequences may be a strong factor in shaping the mutational landscape of SDs. However, while the (A,T) > (C,G) frequency is comparable in SD regions not affected by IGC the mutational landscape at large is still very distinct between SDs and unique parts of the genome. In PCA of the mutational spectra in SDs without IGC, the first principal component captures 94.6% of the variation clearly distinguishing the mutational spectrum of SDs and unique DNA (Fig. S11).

## DISCUSSION

Since the first publications of the human genome (IHGSC 2001; Venter et al. 2001), the pattern of single-nucleotide variation within recently duplicated sequence has been difficult to ascertain, leading initially to the misclassification of PSVs as SNVs (Jeffrey A. Bailey et al. 2002; Fredman et al. 2004). Later, indirect approaches were used to infer true SNVs in SDs but these were far from complete (Nakken et al. 2009). More often than not, large-scale sequencing efforts simply excluded such regions in an effort to prevent PSVs from contaminating SNP databases (1000 Genomes Project Consortium et al. 2015; Zook et al. 2019). The use of phased genome assemblies as opposed to sequence reads had the advantage of allowing us to establish 1:1 orthologous relationships, and the ability to discern the effect of IGC while comparing the pattern of single-nucleotide variation for both duplicated and unique DNA within the same haplotypes. As a result, we identify over 1.99 million nonredundant SNVs in a gene-rich portion of the genome previously considered largely inaccessible.

We find that number of SNVs is elevated by ~60% in duplicated DNA when compared to unique DNA and these findings are consistent with recent primate comparative and *de novo* mutation studies from long-read sequencing data (Logsdon et al. 2021; Noyes et al. 2022), which support an elevated mutation rate for repetitive DNA. We estimate that at least 23% of this increase is due to the action of IGC between paralogous sequences that essentially diversify allelic copies through concerted evolution (Chen et al. 2007). IGC within SDs appears to be remarkably pervasive in the human genome compared to earlier estimates (Dumont and Eichler 2013) with over 32,000 candidate regions (n=799 genes) identified and the average human haplotype showing 1,192 events when compared to the reference. The events can be remarkably large with the top 10% of the size distribution over 14.4 kbp in length having the net effect that entire genes are relocated hundreds of kbp into a new genomic context when compared to the reference. It should be stressed, however, that our method relies on the discovery of a closer match within the reference; by definition this limits the detection of IGC events to regions where the donor sequence is already present in the reference as opposed to an alternate. Moreover, we only interrogated regions where 1:1 synteny could be established. As more of the genome is assessed in the context of a pangenome reference framework, it is likely that the proportion of IGC will increase especially in regions such as the centromere and acrocentric, which currently are not well assembled or characterized (Liao et al. 2022).

One of the most striking features of duplicated DNA is its higher GC content; there is a clear skew in the mutational spectrum of SNVs to maintain this property of SDs beyond expectations from unique DNA. We find a 27.1% increase in transversions that convert cytosine to guanine or the reverse across all triplet contexts. GC-rich DNA has long been regarded as hypermutable. For example, C>G mutations preferentially associate with double-stranded breaks in humans and apes (Jónsson et al. 2017; Rahbari et al. 2016; Goldmann et al. 2018; Gao et al. 2019) and GC-rich regions in yeast show ~2-5 times more mutations depending on sequence context compared to AT-rich DNA (Zhu et al. 2014; Kiktev et al. 2018). Interestingly, in human SD regions, we observe a paucity of CpG transition mutations, characteristically associated with spontaneous deamination of CpG dinucleotides and concomitant transitions (Duncan and Miller 1980). The basis for the latter is unclear but it may be partially explained by the recent observation that duplicated genes show greater degree of hypomethylation when compared to their unique counterparts (Vollger et al. 2022). We propose that excess of guanosine and cytosine transversions is a direct consequence of GC-biased gene conversion (Duret and Galtier 2009) driven by an excess of double-stranded breaks that result from a high rate of NAHR events among paralogous sequences.

## ACKNOWLEDGMENTS

The authors thank T. Brown for help in editing this manuscript.

## Funding

This work was supported in part by grants from the U.S. National Institutes of Health (NIH 5R01HG002385, 5U01HG010971, and 1U01HG010973 to E.E.E., K99HG011041 to P.H., and F31AI150163 to W.S.D.). W.S.D. was supported in part by a Fellowship in Understanding Dynamic and Multi-scale Systems from the James S. McDonnell Foundation. E.E.E. is an investigator of Howard Hughes Medical Institute.

This article is subject to HHMI’s Open Access to Publications policy. HHMI lab heads have previously granted a nonexclusive CC BY 4.0 license to the public and a sublicensable license to HHMI in their research articles. Pursuant to those licenses, the author-accepted manuscript of this article can be made freely available under a CC BY 4.0 license immediately upon publication.

## Author contributions

Conceptualization and design: M.R.V., K.Ha., W.S.D., P.H. and E.E.E. Identification and analysis of SNVs from phased assemblies: M.R.V. Mutational spectrum analysis: M.R.V., W.S.D., M.E.G. and K.H. Evolutionary age analysis: M.R.V. and P.H. Assembly generation: M.A., J.L., B.P. and HPRC. PacBio genome sequence generation: K.M.M., A.P.L., K.Ho. and G.A.L. HPRC Iso-Seq analysis: P.C.D. Copy number analysis and validation: P.C.D., X.G., W.T.H., A.R., D.P. and M.R.V. Table organization: M.R.V. Supplementary material organization: M.R.V. Display items: M.R.V., X.G., P.H. and P.C.D. Resources: HPRC, K.Ha., B.P. and E.E.E. Manuscript writing: M.R.V. and E.E.E. with input from all authors.

## Data and materials availability

PacBio HiFi and ONT data have been deposited into NCBI Sequence Read Archive (SRA) under the following bioproject IDs: PRJNA850430 and PRJNA731524. PacBio HiFi data for CHM1 are under the following SRA accessions: SRX10759865 and SRX10759866. The T2T-CHM13 v1.1 assembly can be found on NCBI (GCA_009914755.3). Code for Snakemake pipelines, data analysis, and figure generation are also available at Zenodo (10.5281/zenodo.6792654) and GitHub (sd-divergence; asm-to-reference-alignment; mutyper_workflow; sd-divergence-and-igc-figures).

## HUMAN PANGENOME REFERENCE CONSORTIUM

Haley J. Abel^1^, Lucinda L Antonacci-Fulton^2^, Mobin Asri^3^, Gunjan Baid^4^, Anastasiya Belyaeva^4^, Konstantinos Billis^5^, Guillaume Bourque^6,7,8^, Silvia Buonaiuto^9^, Andrew Carroll^4^, Mark JP Chaisson^10^, Pi-Chuan Chang^4^, Xian H. Chang^3^, Haoyu Cheng^11,12^, Justin Chu^11^, Sarah Cody^2^, Vincenza Colonna^9,13^, Daniel E. Cook^4^, Omar E. Cornejo^14^, Mark Diekhans^3^, Daniel Doerr^15^, Peter Ebert^15^, Jana Ebler^15^, Evan E. Eichler^16,17^, Jordan M. Eizenga^3^, Susan Fairley^5^, Olivier Fedrigo^18^, Adam L. Felsenfeld^19^, Xiaowen Feng^11,12^, Christian Fischer^20^, Paul Flicek^5^, Giulio Formenti^18^, Adam Frankish^5^, Robert S. Fulton^2^, Yan Gao^21^, Shilpa Garg^22^, Erik Garrison^13^, Carlos Garcia Giron^5^, Richard E. Green^23,24^, Cristian Groza^25^, Andrea Guarracino^26^, Leanne Haggerty^5^, Ira Hall^27,28^, William T Harvey^16^, Marina Haukness^3^, David Haussler^3,17^, Simon Heumos^29,30^, Glenn Hickey^3^, Thibaut Hourlier^5^, Kerstin Howe^31^, Miten Jain^32^, Erich D. Jarvis^33,17^, Hanlee P. Ji^34^, Alexey Kolesnikov^4^, Jan O. Korbel^35^, HoJoon Lee^34^, Alexandra P. Lewis^16^, Heng Li^11,12^, Wen-Wei Liao^2,36,27^, Shuangjia Lu^27^, Tsung-Yu Lu^10^, Julian K. Lucas^3^, Hugo Magalhães^15^, Santiago Marco-Sola^37,38^, Pierre Marijon^15^, Tobias Marschall^15^, Fergal J. Martin^5^, Jennifer McDaniel^39^, Karen H. Miga^3^, Matthew W. Mitchell^40^, Jean Monlong^3^, Jacquelyn Mountcastle^18^, Katherine M. Munson^16^, Moses Njagi Mwaniki^41^, Maria Nattestad^4^, Adam M. Novak^3^, Hugh E. Olsen^3^, Nathan D. Olson^39^, Benedict Paten^3^, Trevor Pesout^3^, Adam M. Phillippy^42^, David Porubsky^16^, Pjotr Prins^13^, Daniela Puiu^43^, Allison A Regier^2^, Arang Rhie^42^, Samuel Sacco^44^, Ashley D. Sanders^45^, Valerie A. Schneider^46^, Baergen I. Schultz^19^, Kishwar Shafin^4^, Jonas A. Sibbesen^47^, Jouni Sirén^3^, Michael W. Smith^19^, Heidi J. Sofia^19^, Ahmad N. Abou Tayoun ^48,49^, Françoise Thibaud-Nissen^46^, Chad Tomlinson^2^, Francesca Floriana Tricomi^5^, Flavia Villani^13^, Mitchell R. Vollger^16,50^, Justin Wagner^39^, Ting Wang^51^, Jonathan M. D. Wood^31^, Aleksey V. Zimin^43,52^, Justin M. Zook^39^

1 Division of Oncology, Department of Internal Medicine, Washington University School of Medicine, St. Louis, MO 63110, USA

2 McDonnell Genome Institute, Washington University School of Medicine, St. Louis, MO 63108, USA

3 UC Santa Cruz Genomics Institute, University of California, Santa Cruz, 1156 High St, Santa Cruz, CA, USA

4 Google LLC, 1600 Amphitheater Pkwy, Mountain View, CA 94043, USA

5 European Molecular Biology Laboratory, European Bioinformatics Institute, Wellcome Genome Campus, Cambridge, CB10 1SD, UK

6 Department of Human Genetics, McGill University, Montreal, Québec H3A 0C7, Canada

7 Canadian Center for Computational Genomics, McGill University, Montreal, Québec H3A 0G1, Canada

8 Institute for the Advanced Study of Human Biology (WPI-ASHBi), Kyoto University, Kyoto 606-8501, Japan

9 Institute of Genetics and Biophysics, National Research Council, Naples 80111, Italy

10 University of Southern California, Quantitative and Computational Biology, 3551 Trousdale, Pkwy, Los Angeles, CA, USA

11 Department of Data Sciences, Dana-Farber Cancer Institute, Boston, MA 02215, USA

12 Department of Biomedical Informatics, Harvard Medical School, Boston, MA 02215, USA

13 Department of Genetics, Genomics and Informatics, University of Tennessee Health Science Center, Memphis, TN 38163, USA

14 School of Biological Sciences, Washington State University, Pullman WA 99163, USA

15 Institute for Medical Biometry and Bioinformatics, Medical Faculty, Heinrich Heine University Düsseldorf, Düsseldorf, Germany

16 Department of Genome Sciences, University of Washington School of Medicine, Seattle, WA 98195, USA

17 Howard Hughes Medical Institute, Chevy Chase, MD 20815, USA

18 The Vertebrate Genome Laboratory, The Rockefeller University, New York, NY 10065, USA

19 National Institutes of Health (NIH)–National Human Genome Research Institute, Bethesda, MD, USA

20 USA University of Tennessee Health Science Center, Memphis, TN 38163, USA

21 Center for Computational and Genomic Medicine, The Children’s Hospital of Philadelphia, Philadelphia, PA 19104, USA.

22 Department of Biology, University of Copenhagen, Denmark

23 Department of Biomolecular Engineering, University of California, Santa Cruz, 1156 High St., Santa Cruz, CA 95064, USA

24 Dovetail Genomics, Scotts Valley, CA 95066, USA

25 Quantitative Life Sciences, McGill University, Montreal, Québec H3A 0C7, Canada

26 Genomics Research Centre, Human Technopole, Milan 20157, Italy

27 Department of Genetics, Yale University School of Medicine, New Haven, CT 06510, USA

28 Center for Genomic Health, Yale University School of Medicine, New Haven, CT 06510, USA

29 Quantitative Biology Center (QBiC), University of Tübingen, Tübingen 72076, Germany

30 Biomedical Data Science, Department of Computer Science, University of Tübingen, Tübingen 72076, Germany

31 Tree of Life, Wellcome Sanger Institute, Hinxton, Cambridge, CB10 1SA, UK

32 Northeastern University, Boston, MA 02115, USA

33 The Rockefeller University, New York, NY 10065, USA

34 Division of Oncology, Department of Medicine, Stanford University School of Medicine, Stanford, CA, 94305, USA

35 European Molecular Biology Laboratory, Genome Biology Unit, Meyerhofstr. 1, 69117 Heidelberg, Germany

36 Department of Medicine, Washington University School of Medicine, St. Louis, MO 63110, USA

37 Computer Sciences Department, Barcelona Supercomputing Center, Barcelona, Spain

38 Departament d’Arquitectura de Computadors i Sistemes Operatius, Universitat Autònoma de Barcelona, Barcelona, Spain

39 Material Measurement Laboratory, National Institute of Standards and Technology, Gaithersburg, MD 20877, USA

40 Coriell Institute for Medical Research, Camden, NJ 08103, USA

41 Department of Computer Science, University of Pisa, Pisa 56127, Italy

42 Genome Informatics Section, Computational and Statistical Genomics Branch, National Human Genome Research Institute, National Institutes of Health, Bethesda, MD 20892, USA

43 Department of Biomedical Engineering, Johns Hopkins University, Baltimore 21218, MD, USA

44 Department of Ecology & Evolutionary Biology, University of California, Santa Cruz, 1156 High St, Santa Cruz, CA, USA

45 Berlin Institute for Medical Systems Biology, Max Delbrück Center for Molecular Medicine in the Helmholtz Association, Berlin, Germany

46 National Center for Biotechnology Information, National Library of Medicine, National Institutes of Health, Bethesda, MD 20894, USA

47 Center for Health Data Science, University of Copenhagen, Denmark

48 Al Jalila Genomics Center of Excellence, Al Jalila Children’s Specialty Hospital, Dubai, UAE

49 Center for Genomic Discovery, Mohammed Bin Rashid University of Medicine and Health Sciences, Dubai, UAE

50 Division of Medical Genetics, University of Washington School of Medicine, Seattle, WA 98195, USA

51 Department of Genetics, Washington University School of Medicine, St. Louis, MO 63110, USA

52 Center for Computational Biology, Johns Hopkins University, Baltimore, MD 21218, USA

## References

1000 Genomes Project Consortium, Gonçalo R. Abecasis, David Altshuler, Adam Auton, Lisa D. Brooks, Richard M. Durbin, Richard A. Gibbs, Matt E. Hurles, and Gil A. McVean. 2010. “A Map of Human Genome Variation from Population-Scale Sequencing.” Nature 467 (7319): 1061–73. https://doi.org/10.1038/nature09534.

1000 Genomes Project Consortium, Goncalo R. Abecasis, Adam Auton, Lisa D. Brooks, Mark A. DePristo, Richard M. Durbin, Robert E. Handsaker, Hyun Min Kang, Gabor T. Marth, and Gil A. McVean. 2012. “An Integrated Map of Genetic Variation from 1,092 Human Genomes.” Nature 491 (7422): 56–65. https://doi.org/10.1038/nature11632.

1000 Genomes Project Consortium, Adam Auton, Lisa D. Brooks, Richard M. Durbin, Erik P. Garrison, Hyun Min Kang, Jan O. Korbel, et al. 2015. “A Global Reference for Human Genetic Variation.” Nature 526 (7571): 68–74. https://doi.org/10.1038/nature15393.

Amemiya, Haley M., Anshul Kundaje, and Alan P. Boyle. 2019. “The ENCODE Blacklist: Identification of Problematic Regions of the Genome.” Scientific Reports 9 (1): 9354. https://doi.org/10.1038/s41598-019-45839-z.

Bailey, J. A., A. M. Yavor, H. F. Massa, B. J. Trask, and E. E. Eichler. 2001. “Segmental Duplications: Organization and Impact within the Current Human Genome Project Assembly.” Genome Research 11 (6): 1005–17. https://doi.org/10.1101/gr.gr-1871r.

Bailey, Jeffrey A., Zhiping Gu, Royden A. Clark, Knut Reinert, Rhea V. Samonte, Stuart Schwartz, Mark D. Adams, Eugene W. Myers, Peter W. Li, and Evan E. Eichler. 2002. “Recent Segmental Duplications in the Human Genome.” Science 297 (5583): 1003–7. https://doi.org/10.1126/science.1072047.

Bailey, Jeffrey A., Ge Liu, and Evan E. Eichler. 2003. “An Alu Transposition Model for the Origin and Expansion of Human Segmental Duplications.” American Journal of Human Genetics 73 (4): 823–34. https://doi.org/10.1086/378594.

Bosch, Elena, Matthew E. Hurles, Arcadi Navarro, and Mark A. Jobling. 2004. “Dynamics of a Human Interparalog Gene Conversion Hotspot.” Genome Research. https://doi.org/10.1101/gr.2177404.

Carlson, Jedidiah, William S. DeWitt, and Kelley Harris. 2020. “Inferring Evolutionary Dynamics of Mutation Rates through the Lens of Mutation Spectrum Variation.” Current Opinion in Genetics & Development 62 (June): 50–57. https://doi.org/10.1016/j.gde.2020.05.024.

Chaisson, Mark J. P., John Huddleston, Megan Y. Dennis, Peter H. Sudmant, Maika Malig, Fereydoun Hormozdiari, Francesca Antonacci, et al. 2015. “Resolving the Complexity of the Human Genome Using Single-Molecule Sequencing.” Nature 517 (7536): 608–11. https://doi.org/10.1038/nature13907.

Chaisson, Mark J. P., Ashley D. Sanders, Xuefang Zhao, Ankit Malhotra, David Porubsky, Tobias Rausch, Eugene J. Gardner, et al. 2019. “Multi-Platform Discovery of Haplotype-Resolved Structural Variation in Human Genomes.” Nature Communications 10 (1): 1784. https://doi.org/10.1038/s41467-018-08148-z.

Cheng, Haoyu, Gregory T. Concepcion, Xiaowen Feng, Haowen Zhang, and Heng Li. 2021. “Haplotype-Resolved de Novo Assembly Using Phased Assembly Graphs with Hifiasm.” Nature Methods 18 (2): 170–75. https://doi.org/10.1038/s41592-020-01056-5.

Chen, Jian-Min, David N. Cooper, Nadia Chuzhanova, Claude Férec, and George P. Patrinos. 2007. “Gene Conversion: Mechanisms, Evolution and Human Disease.” Nature Reviews. Genetics 8 (10): 762–75. https://doi.org/10.1038/nrg2193.

Conant, Gavin C., and Andreas Wagner. 2003. “Asymmetric Sequence Divergence of Duplicate Genes.” Genome Research 13 (9): 2052–58. https://doi.org/10.1101/gr.1252603.

DeWitt, William S. 2020. “Mutyper: Assigning and Summarizing Mutation Types for Analyzing Germline Mutation Spectra.” bioRxiv. https://doi.org/10.1101/2020.07.01.183392.

Dishuck, Philip C., Allison N. Rozanski, Glennis A. Logsdon, and Evan E. Eichler. 2022. “GAVISUNK: Genome Assembly Validation via Inter-SUNK Distances in Oxford Nanopore Reads.” bioRxiv. https://doi.org/10.1101/2022.06.17.496619.

Dumont, Beth L., and Evan E. Eichler. 2013. “Signals of Historical Interlocus Gene Conversion in Human Segmental Duplications.” PloS One 8 (10): e75949. https://doi.org/10.1371/journal.pone.0075949.

Duncan, Bruce K., and Jeffrey H. Miller. 1980. “Mutagenic Deamination of Cytosine Residues in DNA.” Nature. https://doi.org/10.1038/287560a0.

Duret, Laurent, and Nicolas Galtier. 2009. “Biased Gene Conversion and the Evolution of Mammalian Genomic Landscapes.” Annual Review of Genomics and Human Genetics 10 (1): 285–311. https://doi.org/10.1146/annurev-genom-082908-150001.

Ebert, Peter, Peter A. Audano, Qihui Zhu, Bernardo Rodriguez-Martin, David Porubsky, Marc Jan Bonder, Arvis Sulovari, et al. 2021. “Haplotype-Resolved Diverse Human Genomes and Integrated Analysis of Structural Variation.” Science 372 (6537). https://doi.org/10.1126/science.abf7117.

Ebler, Jana, Peter Ebert, Wayne E. Clarke, Tobias Rausch, Peter A. Audano, Torsten Houwaart, Yafei Mao, et al. 2022. “Pangenome-Based Genome Inference Allows Efficient and Accurate Genotyping across a Wide Spectrum of Variant Classes.” Nature Genetics 54 (4): 518–25. https://doi.org/10.1038/s41588-022-01043-w.

Elliott, B., C. Richardson, J. Winderbaum, J. A. Nickoloff, and M. Jasin. 1998. “Gene Conversion Tracts from Double-Strand Break Repair in Mammalian Cells.” Molecular and Cellular Biology 18 (1): 93–101. https://doi.org/10.1128/MCB.18.1.93.

Force, A., M. Lynch, F. B. Pickett, A. Amores, Y. L. Yan, and J. Postlethwait. 1999. “Preservation of Duplicate Genes by Complementary, Degenerative Mutations.” Genetics 151 (4): 1531–45. https://www.ncbi.nlm.nih.gov/pubmed/10101175.

Fredman, David, Stefan J. White, Susanna Potter, Evan E. Eichler, Johan T. Den Dunnen, and Anthony J. Brookes. 2004. “Complex SNP-Related Sequence Variation in Segmental Genome Duplications.” Nature Genetics 36 (8): 861–66. https://doi.org/10.1038/ng1401.

Gao, Ziyue, Priya Moorjani, Thomas A. Sasani, Brent S. Pedersen, Aaron R. Quinlan, Lynn B. Jorde, Guy Amster, and Molly Przeworski. 2019. “Overlooked Roles of DNA Damage and Maternal Age in Generating Human Germline Mutations.” Proceedings of the National Academy of Sciences of the United States of America 116 (19): 9491–9500. https://doi.org/10.1073/pnas.1901259116.

Goldmann, Jakob M., Vladimir B. Seplyarskiy, Wendy S. W. Wong, Thierry Vilboux, Pieter B. Neerincx, Dale L. Bodian, Benjamin D. Solomon, et al. 2018. “Germline de Novo Mutation Clusters Arise during Oocyte Aging in Genomic Regions with High Double-Strand-Break Incidence.” Nature Genetics 50 (4): 487–92. https://doi.org/10.1038/s41588-018-0071-6.

Harris, Kelley. 2015. “Evidence for Recent, Population-Specific Evolution of the Human Mutation Rate.” Proceedings of the National Academy of Sciences of the United States of America 112 (11): 3439–44. https://doi.org/10.1073/pnas.1418652112.

Hsieh, Pinghsun, Vy Dang, Mitchell Vollger, Yafei Mao, Tzu-Hsueh Huang, Philip Dishuck, Carl Baker, et al. 2020. “Opposing Selective Forces Operating on Human-Specific Duplicated TCAF Genes in Neanderthals and Humans.”

Huddleston, John, Mark J. P. Chaisson, Karyn Meltz Steinberg, Wes Warren, Kendra Hoekzema, David Gordon, Tina A. Graves-Lindsay, et al. 2017. “Discovery and Genotyping of Structural Variation from Long-Read Haploid Genome Sequence Data.” Genome Research 27 (5): 677–85. https://doi.org/10.1101/gr.214007.116.

Hurles, Matthew. 2002. “Are 100,000 ‘SNPs’ Useless?” Science. American Association for the Advancement of Science (AAAS). https://doi.org/10.1126/science.298.5598.1509a.

IHGSC. 2001. “Initial Sequencing and Analysis of the Human Genome.” Nature 409 (6822): 860–921. https://doi.org/10.1038/35057062.

Jarvis, Erich D., Giulio Formenti, Arang Rhie, Andrea Guarracino, Chentao Yang, Jonathan Wood, Alan Tracey, et al. 2022. “Automated Assembly of High-Quality Diploid Human Reference Genomes.” bioRxiv. https://doi.org/10.1101/2022.03.06.483034.

Jiang, Z., H. Tang, M. Ventura, M. F. Cardone, T. Marques-Bonet, X. She, P. a. Pevzner, and E. E. Eichler. 2007. “Ancestral Reconstruction of Segmental Duplications Reveals Punctuated Cores of Human Genome Evolution.” Nature Genetics 39 (11): 1361–68. https://doi.org/10.1038/ng.2007.9.

Jónsson, Hákon, Patrick Sulem, Birte Kehr, Snaedis Kristmundsdottir, Florian Zink, Eirikur Hjartarson, Marteinn T. Hardarson, et al. 2017. “Parental Influence on Human Germline de Novo Mutations in 1,548 Trios from Iceland.” Nature 549 (7673): 519–22. https://doi.org/10.1038/nature24018.

Kiktev, Denis A., Ziwei Sheng, Kirill S. Lobachev, and Thomas D. Petes. 2018. “GC Content Elevates Mutation and Recombination Rates in the Yeast Saccharomyces Cerevisiae.” Proceedings of the National Academy of Sciences of the United States of America 115 (30): E7109–18. https://doi.org/10.1073/pnas.1807334115.

Lek, Monkol, Konrad J. Karczewski, Eric V. Minikel, Kaitlin E. Samocha, Eric Banks, Timothy Fennell, Anne H. O'Donnell-Luria, et al. 2016. “Analysis of Protein-Coding Genetic Variation in 60,706 Humans.” Nature 536 (7616): 285–91. https://doi.org/10.1038/nature19057.

Liao, Wen-Wei, Mobin Asri, Jana Ebler, Daniel Doerr, Marina Haukness, Glenn Hickey, Shuangjia Lu, et al. 2022. “A Draft Human Pangenome Reference.” bioRxiv, July.

Logsdon, Glennis A., Mitchell R. Vollger, Pinghsun Hsieh, Yafei Mao, Mikhail A. Liskovykh, Sergey Koren, Sergey Nurk, et al. 2021. “The Structure, Function and Evolution of a Complete Human Chromosome 8.” Nature 593 (7857): 101–7. https://doi.org/10.1038/s41586-021-03420-7.

McCarroll, Steven A., Tracy N. Hadnott, George H. Perry, Pardis C. Sabeti, Michael C. Zody, Jeffrey C. Barrett, Stephanie Dallaire, et al. 2006. “Common Deletion Polymorphisms in the Human Genome.” Nature Genetics 38 (1): 86–92. https://doi.org/10.1038/ng1696.

Nakken, Sigve, Einar A. Rødland, Torbjørn Rognes, and Eivind Hovig. 2009. “Large-Scale Inference of the Point Mutational Spectrum in Human Segmental Duplications.” BMC Genomics 10 (January): 43. https://doi.org/10.1186/1471-2164-10-43.

Noyes, Michelle D., William T. Harvey, David Porubsky, Arvis Sulovari, Ruiyang Li, Nicholas R. Rose, Peter A. Audano, et al. 2022. “Familial Long-Read Sequencing Increases Yield of de Novo Mutations.” American Journal of Human Genetics 109 (4): 631–46. https://doi.org/10.1016/j.ajhg.2022.02.014.

Nurk, Sergey, Sergey Koren, Arang Rhie, Mikko Rautiainen, Andrey V. Bzikadze, Alla Mikheenko, Mitchell R. Vollger, et al. 2022. “The Complete Sequence of a Human Genome.” Science 376 (6588): 44–53. https://doi.org/10.1126/science.abj6987.

Pietri, Mathéa, Caroline Dakowski, Samia Hannaoui, Aurélie Alleaume-Butaux, Julia Hernandez-Rapp, Audrey Ragagnin, Sophie Mouillet-Richard, et al. 2013. “PDK1 Decreases TACE-Mediated α-Secretase Activity and Promotes Disease Progression in Prion and Alzheimer's Diseases.” Nature Medicine 19 (9): 1124–31. https://doi.org/10.1038/nm.3302.

Rahbari, Raheleh, UK10K Consortium, Arthur Wuster, Sarah J. Lindsay, Robert J. Hardwick, Ludmil B. Alexandrov, Saeed Al Turki, et al. 2016. “Timing, Rates and Spectra of Human Germline Mutation.” Nature Genetics. https://doi.org/10.1038/ng.3469.

Rautiainen, Mikko, Sergey Nurk, Brian P. Walenz, Glennis A. Logsdon, David Porubsky, Arang Rhie, Evan E. Eichler, Adam M. Phillippy, and Sergey Koren. 2022. “Verkko: Telomere-to-Telomere Assembly of Diploid Chromosomes.” bioRxiv. https://doi.org/10.1101/2022.06.24.497523.

Richter, Melanie, Nadeem Murtaza, Robin Scharrenberg, Sean H. White, Ole Johanns, Susan Walker, Ryan K. C. Yuen, et al. 2019. “Altered TAOK2 Activity Causes Autism-Related Neurodevelopmental and Cognitive Abnormalities through RhoA Signaling.” Molecular Psychiatry 24 (9): 1329–50. https://doi.org/10.1038/s41380-018-0025-5.

Sawyer. n.d. “GENECONV: A Computer Package for the Statistical Detection of Gene Conversion.” Http://www.Math.Wustl.Edu/~Sawyer. https://ci.nii.ac.jp/naid/10027221513/.

Sawyer, S. 1989. “Statistical Tests for Detecting Gene Conversion.” Molecular Biology and Evolution 6 (5): 526–38. https://doi.org/10.1093/oxfordjournals.molbev.a040567.

Schneider, Valerie A., Tina Graves-Lindsay, Kerstin Howe, Nathan Bouk, Hsiu Chuan Chen, Paul A. Kitts, Terence D. Murphy, et al. 2017. “Evaluation of GRCh38 and de Novo Haploid Genome Assemblies Demonstrates the Enduring Quality of the Reference Assembly.” Genome Research 27 (5). https://doi.org/10.1101/gr.213611.116.

Sekar, Aswin, Allison R. Bialas, Heather de Rivera, Avery Davis, Timothy R. Hammond, Nolan Kamitaki, Katherine Tooley, et al. 2016. “Schizophrenia Risk from Complex Variation of Complement Component 4.” Nature 530 (7589): 177–83. https://doi.org/10.1038/nature16549.

Shahi, Abbas, Shima Afzali, Saeedeh Salehi, Saeed Aslani, Mahdi Mahmoudi, Ahmadreza Jamshidi, and Aliakbar Amirzargar. 2020. “IL-27 and Autoimmune Rheumatologic Diseases: The Good, the Bad, and the Ugly.” International Immunopharmacology 84 (July): 106538. https://doi.org/10.1016/j.intimp.2020.106538.

Sudmant, Peter H., Jacob O. Kitzman, Francesca Antonacci, Can Alkan, Maika Malig, Anya Tsalenko, Nick Sampas, Laurakay Bruhn, and Jay Shendure. 2010. “Diversity of Human Copy Number.” Science 11184 (2009): 2–7. https://doi.org/papers2://publication/uuid/C37D7A2A-43D1-4164-A5DE-4447C58EBC0D.

Sudmant, Peter H., Tobias Rausch, Eugene J. Gardner, Robert E. Handsaker, Alexej Abyzov, John Huddleston, Yan Zhang, et al. 2015. “An Integrated Map of Structural Variation in 2,504 Human Genomes.” Nature 526 (7571): 75–81. https://doi.org/10.1038/nature15394.

Teshima, Kosuke M., and Hideki Innan. 2012. “The Coalescent with Selection on Copy Number Variants.” Genetics 190 (3): 1077–86. https://doi.org/10.1534/genetics.111.135343.

Venter, J. C., M. D. Adams, E. W. Myers, P. W. Li, R. J. Mural, G. G. Sutton, H. O. Smith, et al. 2001. “The Sequence of the Human Genome.” Science 291 (5507): 1304–51. https://doi.org/10.1126/science.1058040.

Vollger, Mitchell R., Xavi Guitart, Philip C. Dishuck, Ludovica Mercuri, William T. Harvey, Ariel Gershman, Mark Diekhans, et al. 2022. “Segmental Duplications and Their Variation in a Complete Human Genome.” Science 376 (6588): eabj6965. https://doi.org/10.1126/science.abj6965.

Vollger, Mitchell R., Glennis A. Logsdon, Peter A. Audano, Arvis Sulovari, David Porubsky, Paul Peluso, Aaron M. Wenger, et al. 2020. “Improved Assembly and Variant Detection of a Haploid Human Genome Using Single-Molecule, High-Fidelity Long Reads.” Annals of Human Genetics 84 (2): 125–40. https://doi.org/10.1111/ahg.12364.

Zhang, Ze, Z. W. Luo, Hirohisa Kishino, and Mike J. Kearsey. 2005. “Divergence Pattern of Duplicate Genes in Protein-Protein Interactions Follows the Power Law.” Molecular Biology and Evolution 22 (3): 501–5. https://doi.org/10.1093/molbev/msi034.

Zhu, Yuan O., Mark L. Siegal, David W. Hall, and Dmitri A. Petrov. 2014. “Precise Estimates of Mutation Rate and Spectrum in Yeast.” Proceedings of the National Academy of Sciences of the United States of America 111 (22): E2310–18. https://doi.org/10.1073/pnas.1323011111.

Zook, Justin M., Jennifer McDaniel, Nathan D. Olson, Justin Wagner, Hemang Parikh, Haynes Heaton, Sean A. Irvine, et al. 2019. “An Open Resource for Accurately Benchmarking Small Variant and Reference Calls.” Nature Biotechnology 37 (5): 561–66. https://doi.org/10.1038/s41587-019-0074-6.

