## Supplement for "Increased mutation rate and interlocus gene conversion within human segmental duplications"

### Supplemental methods and information for: Increased mutation rate and interlocus gene conversion within human segmental duplications.

#### Supplemental Information

**Quality of HPRC segmental duplication (SD) content.** Compared to previous genome assemblies focused on human diversity (Ebert *et al.*, 2021), Human Pangenome Reference Consortium (HPRC) assemblies of SD regions are significantly more contiguous (Liao *et al.*, 2022). However, SDs are frequently a source misassembly, so we performed a number of validation experiments to further assess the quality of HPRC SDs. Using the T2T reference as a guide of completeness and the HPRC callset of potentially unreliable regions (Liao *et al.*, 2022), we determined that on average only 1.64 Mbp (1.37%) of the analyzed SD sequence was suspect due to abnormal read coverage.

First, we selected 19 copy number variable duplicated loci and compared Illumina WGS read-depth estimates to that predicted by k-merized counts based on summing the two haplotypes from each of the 47 individuals present in the HPRC phase one assemblies (Methods). We observe a striking correlation between the two (r2=0.97) with 756 of 893 tests predicting the exact same copy number (Figure S1) suggesting that the vast majority of SD gene structures are haplotype resolved with appropriate copy number within HPRC samples. Over half of the failed copy number estimate comparisons occurred within SDs that span greater than 141 kbp and which are more than 99.3% identical on average. These constitute some of the largest and most identical SDs in the human genome where extensive structural variation exists.

As a second test for haplotype integrity, we aligned orthogonal Oxford Nanopore Technologies (ONT) sequencing data produced from the same samples. While less accurate than HiFi data, ONT data has the advantage that it is on average three times longer allowing large repetitive regions to be effectively spanned and scaffolded (Logsdon *et al.*, 2021). We devised a strategy to phase (Koren *et al.*, 2018) and align reads based on matching singly unique nucleotide k-mers (SUNKs) and their distance between the assemblies and the ONT data (Dishuck *et al.*, 2022). We applied this method to assemblies with ultra-long ONT data available (n=35) and, in particular, to IGC acceptor tracts. On average, 94% of acceptor tracts validate by ONT inter-SUNK distances, with an interquartile distance of 94-98%. For 52 large duplicated loci, we found that 95.9% of the base pairs validate by inter-SUNK distances in ONT reads. While gaps remain and are preferentially enriched in regions of the largest and most identical SDs (see Porubsky et al., in preparation), these analyses confirm haplotype contiguity allowing for patterns of SNV variation to be assessed for the first time.

As a final control for potential haplotype mixing of long reads from diploid samples confounding SNV diversity analyses, we generated deep ONT and HiFi data from a second hydatidiform mole where a single paternal haplotype was present. Using an alternate assembler (Verkko) that leverages both HiFi and ONT data, we generated a highly contiguous (contig N50 = 110.90 Mbp) and highly accurate (QV=55) assembly of another haploid human genome (Rautiainen *et al.*, 2022). SDs were predictably more contiguous in this haploid sample when compared to the HPRC assemblies though the difference was modest. In the CHM1 assembly we identify an additional 11.9 Mbp of 1-1 sequence alignment (total of 132 Mbp) for SDs when compared to the HPRC hifiasm assemblies (132,082,329-120,190,200 bp, Figure 1a). We used this second hydatidiform mole as an internal control for all analyses focused on SNV diversity and mutation rate analyses (below). Our analysis showed a more complete assembly of CHM1 did not impact our observations of increased mutational rate across any category. CHM1 has, on average, 12.55 SNVs per 10 kbp, which is in agreement with all the other non-African samples (Figure 1d). Furthermore, the Mbp of interlocus gene conversion (IGC) as well as the total number of events in CHM1 (6.69 Mbp, 1,118 events) is very comparable to the other haplotype assemblies [7.47 Mbp, 1,192 events (6.04 Mbp in Europeans)].

**TCAF geneconv validation.** To assess the performance of our IGC-calling method, we chose to apply an alternative procedure using a model-based approach implemented in the program GENECONV (v.1.81a) for IGC detection (Sawyer). Briefly, GENECONV identifies pairs of uninterrupted sequences with nearly 100% sequence identity that are longer than expected given the overall pattern of variable sites in an alignment. We first identify paralogous sequences and align them using MAFFT (v7.453; mafft --maxiterate 100 sequences.fasta > aln.fasta) and run GENECONV using the following command: geneconv aln.fasta -Annotate -Minnpoly=1 -nolog /lp. To be conservative, we only considered IGC tracts that have both simulation- and Karlin-Altschul- p values < 0.05 reported by GENECONV. We applied this approach to re-examine the *TCAF* locus on chromosome 7q35, which has been shown to have wide-spread IGC events in the human genome (Hsieh *et al.* 2021). In total, GENECONV identifies 87 potential IGC events. Of note, 15 out of 18 highly supported IGC loci (at least 20 supporting SNVs) identified by our alignment-based method are also identified in the 87 GENECONV-based IGC loci. This result indicates that our method for IGC detection is more conservative and highly specific.

### Supplemental Methods

**Copy number estimate validation.** The goal of this analysis was to validate copy number from the assembled HPRC haplotypes compared to estimates from read-depth analysis of the same samples sequenced using Illumina WGS. Large, recently duplicated segments are prone to copy number variation and are also susceptible to collapse and misassembly due to their repetitive nature. HPRC haplotypes were assembled using circular consensus sequence or CCS with hifiasm (Cheng *et al.*, 2021; Liao *et al.*, 2022) creating contiguous long-read assemblies. We selected 19 SD loci corresponding to genes that were known to be duplicated and copy number variable in the human species. We k-merized the two haplotype assemblies corresponding to each locus for each individual into k-mers of 31 base pairs in length. We then computed copy number estimates over each locus for the sum haplotype assemblies and calculated the difference based on Illumina WGS from the same sample. For both datasets we derived these estimates using FastCN, an algorithm implementing whole-genome shotgun sequence detection (Pendleton et al. 2018). When averaging across each region and comparing differences in assembly copy versus Illumina WGS copy estimate, we observe that 756 out of 893 tests were perfectly matched (delta=0), suggesting the majority of these assemblies correctly represent the underlying genomic sequence of the samples.

**Haplotype integrity analysis using inter-SUNK approach.** For the 35 HPRC assemblies with matched ultra-long ONT data, we applied the GAVISUNK method as an orthogonal validation of HiFi assembly integrity (https://github.com/pdishuck/GAVISUNK, (Dishuck *et al.*, 2022). Briefly, candidate haplotype-specific SUNKs are determined from the HiFi assembly and compared to ONT reads phased with parental Illumina data. Inter-SUNK distances are required to be consistent between the assembly and ONT reads, and regions that can be spanned and tiled with consistent ONT reads are considered validated. ONT read dropouts do not necessarily correspond to misassembly—they are also caused by large regions devoid of haplotype-specific SUNKs from recent duplications, homozygosity, or overassembly of the region, as well as Poisson dropout of read coverage.

**Read-depth analysis using the HPRC unreliable callset.** For the 94 assembled HPRC haplotypes, we downloaded the regions identified to have abnormal coverage using form S3 (s3://human-pangenomics/submissions/e9ad8022-1b30-11ec-ab04-0a13c5208311--COVERAGE_ANALYSIS_Y1_GENBANK/FLAGGER/JAN_09_2022/FINAL_HIFI_BASED/FLAGGER_HIFI_ASM_SIMPLIFIED_BEDS/ALL/). We then intersected these regions with the callable SD regions within each assembly to determine the number of collapsed, falsely duplicated, and low-coverage base pairs within each assembly. The unreliable regions were determined by the HPRC using the Flagger pipeline (https://github.com/human-pangenomics/hpp_production_workflows/tree/asset/coverage).

**Whole-genome alignments and synteny definition.** Whole-genome alignments were calculated against T2T-CHM13 v1.1 with a copy of GRCh38 chrY using minimap2 v2.24 with these parameters "-a -x asm20 --secondary=no -s 25000 -K 8G". The alignments were further processed with rustybam v0.1.29 using the subcommands trim-paf, to remove alignments that were redundant in query sequence, and break-paf, to split alignments on structural variants over 10 kbp. After these steps the remaining alignments over 1 Mbp of continuously alignmed sequence were defined to be syntenic. The pipeline is available on github at: https://github.com/mrvollger/asm-to-reference-alignment/.

**Estimating the diversity of SNVs in SDs and unique sequences.** When enumerating the number of SNVs, we count all pairwise differences between the haplotypes and the reference, counting events observed in multiple haplotypes multiple times. Therefore, except where otherwise indicated we are referring to the total number of pairwise differences rather than the total number of nonredundant SNVs (number of segregation sites).

**Defining IGC events.** The query sequence with syntenic alignments was fragmented into 1 kbp windows with a 100 bp slide and realigned back to T2T-CHM13 v1.1 using minimap2 v2.24 to identify each window's single best alignment position. These alignments were compared to their original syntenic alignment positions, and if they were not overlapping, we considered them to be candidate IGC windows. Candidate IGC windows were then merged into larger intervals when windows were overlapping in both the donor and the acceptor sequence. These merged candidate IGC windows were once again aligned to determine the full extent of the IGC event in a single alignment. If the IGC event was still nonoverlapping with the syntenic alignment at this stage, we used the CIGAR string to identify the number of matching and mismatching bases at the "donor" site and compared that to the number of matching and mismatching bases at the acceptor site determined by the syntenic alignment. Code available on github: https://github.com/mrvollger/asm-to-reference-alignment/.

**Chimpanzee genome assembly.** We assembled the genome of the now deceased chimpanzee sample Clint (S006007) using HiFi long-read data and hifiasm version 0.15.2. The assembly is locally phased since trio-binning and HiC data were unavailable. Data is available on NCBI SRA under the bioproject PRJNA659034 (Logsdon *et al.*, 2021).

**Determining the composition of triplet mutations in SD and unique sequences:** Mutational spectra for unique and SD regions from each individual were computed using mutyper (DeWitt 2020; Carlson, DeWitt, and Harris 2020) based on derived SNVs polarized against a chimpanzee genome assembly as described above. These spectra were normalized to the triplet content of the respective unique or SD regions by dividing the count of each triplet mutation type by the total count of each triplet context in the ancestral region and normalizing the number of counts in SD and unique sequences to be the same. For PCA the data was further normalized using the centered log-ratio transformation which is commonly used for compositional measurements (Aitchison, 1982).

**Estimation of time to the most recent common ancestor (TMRCA)**

To estimate TMRCA for a locus of interest, we focus on orthologous sequences (10 kbp windows) identified in synteny among human and chimpanzee haplotypes. Under an assumption of infinite sites, the number of mutations $x_i$ between a human sequence and its MRCA is Poisson distributed with a mean of $\mu \times T$, where $\mu$ is the mutation rate scaled with respect to the substitutions between human and chimpanzee lineages, and T is the TMRCA. That is, $T = \sum_{i=1}^{n} x_i / n\mu$, where n is the number of human haplotypes. To convert TMRCA to time in years, we assume six million years of divergence between human and chimpanzee lineages.

### Supplemental Figures

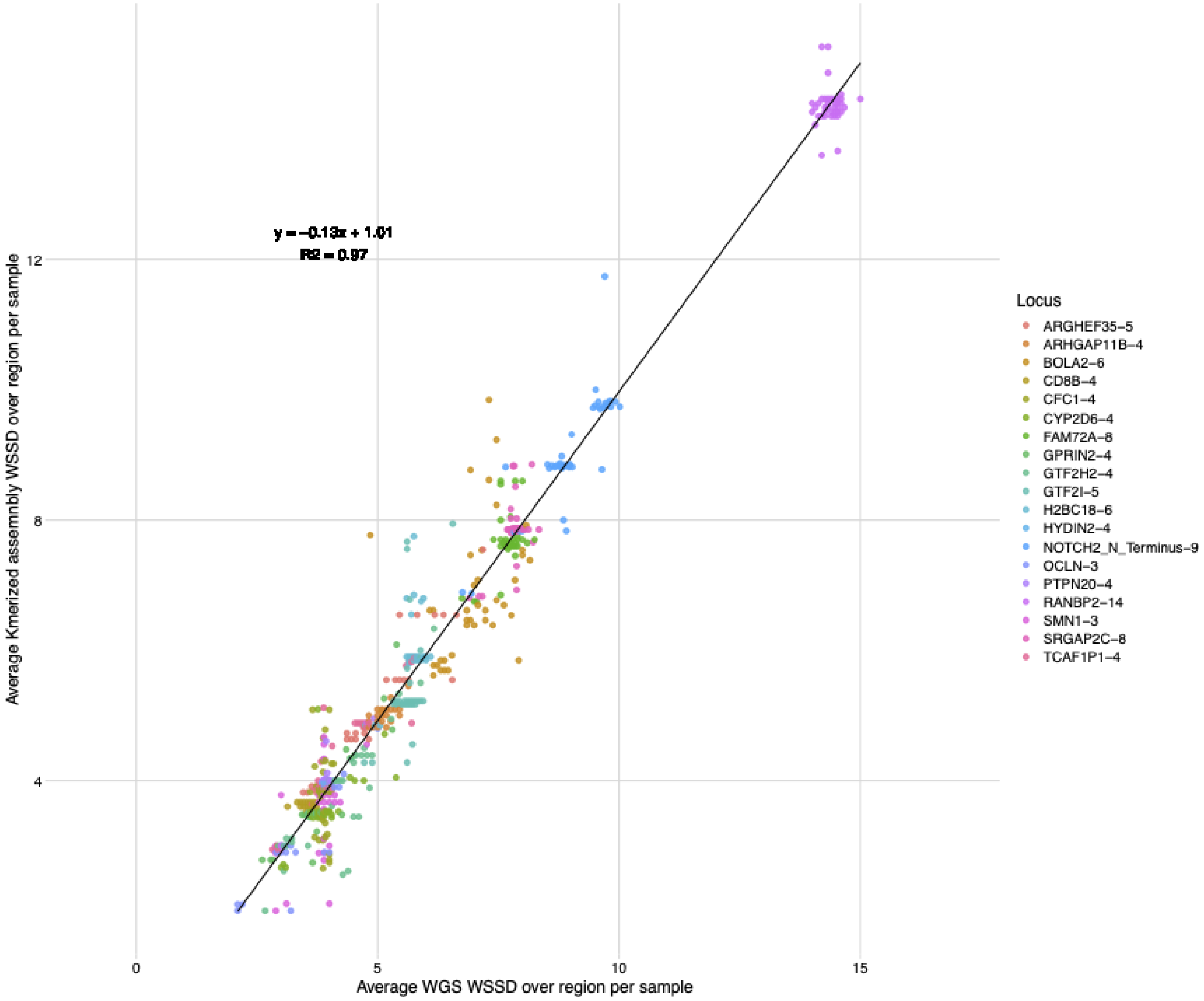


**Figure S1. Copy number estimate correlation.** Here we illustrate the correlation between each sample's diplotype assembly copy number estimate and the corresponding NGS WSSD copy number estimates from libraries generated in the 1000 Genomes Project for 19 selected SD loci. We observe a strong correlation between the two estimates (r2 = 0.97), indicating that, by copy number, the assemblies generated by the HPRC support native genomic copy number in these SD regions.

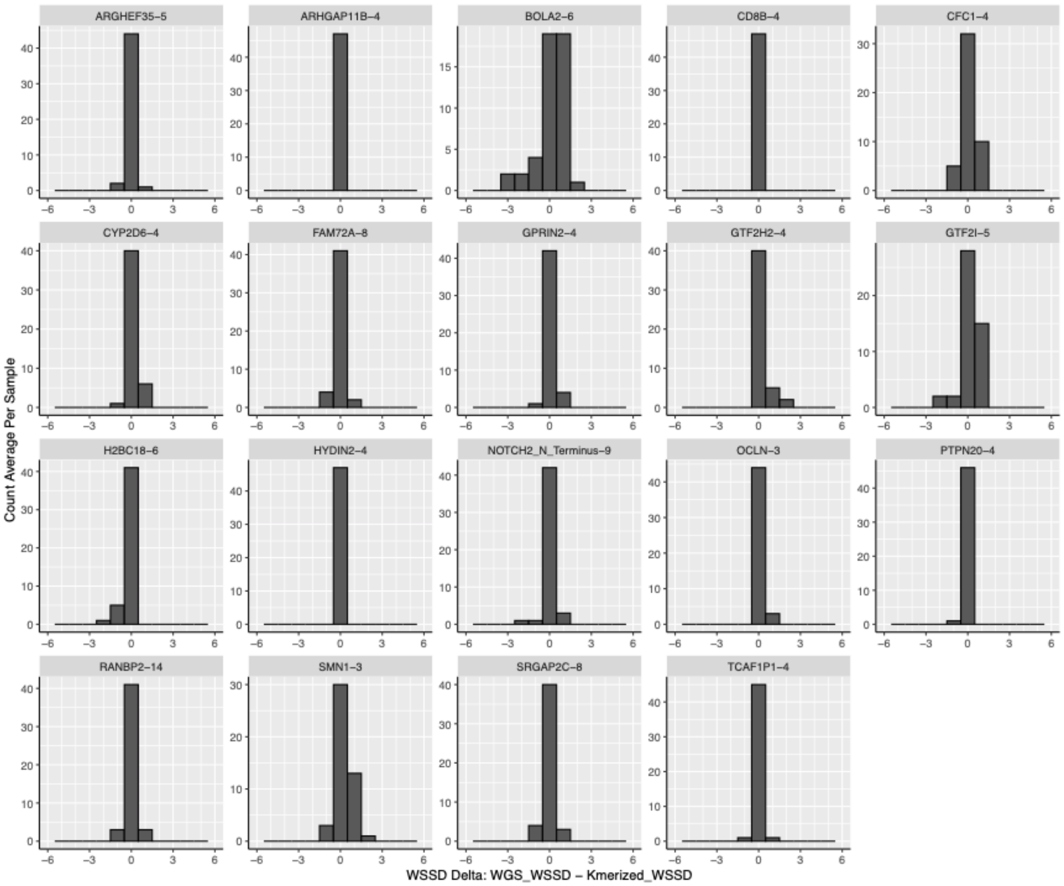


**Figure S2. Average copy number estimate comparison.** We estimated the copy number of 19 human SDs across all 47 samples (96 haplotypes) using either k-mers aggregated from both assembly haplotypes or orthogonal NGS developed by the 1000 Genomes Project. Here we illustrate the average copy number estimate differences over these 19 regions between the two methods. Titles of each histogram correspond to the gene models residing within, and commonly associate with the particular HSD region selected. Each number trailing the gene models are median copy number estimates for these regions across all 47 samples, estimated by diplotype assembly copy number. For the majority of tests, 756 of 893, copy number estimates are identical, resulting in a delta of zero. Diverging deltas most commonly occur in the largest and most highly identical regions but may also diverge due to region size or lower identity duplications.

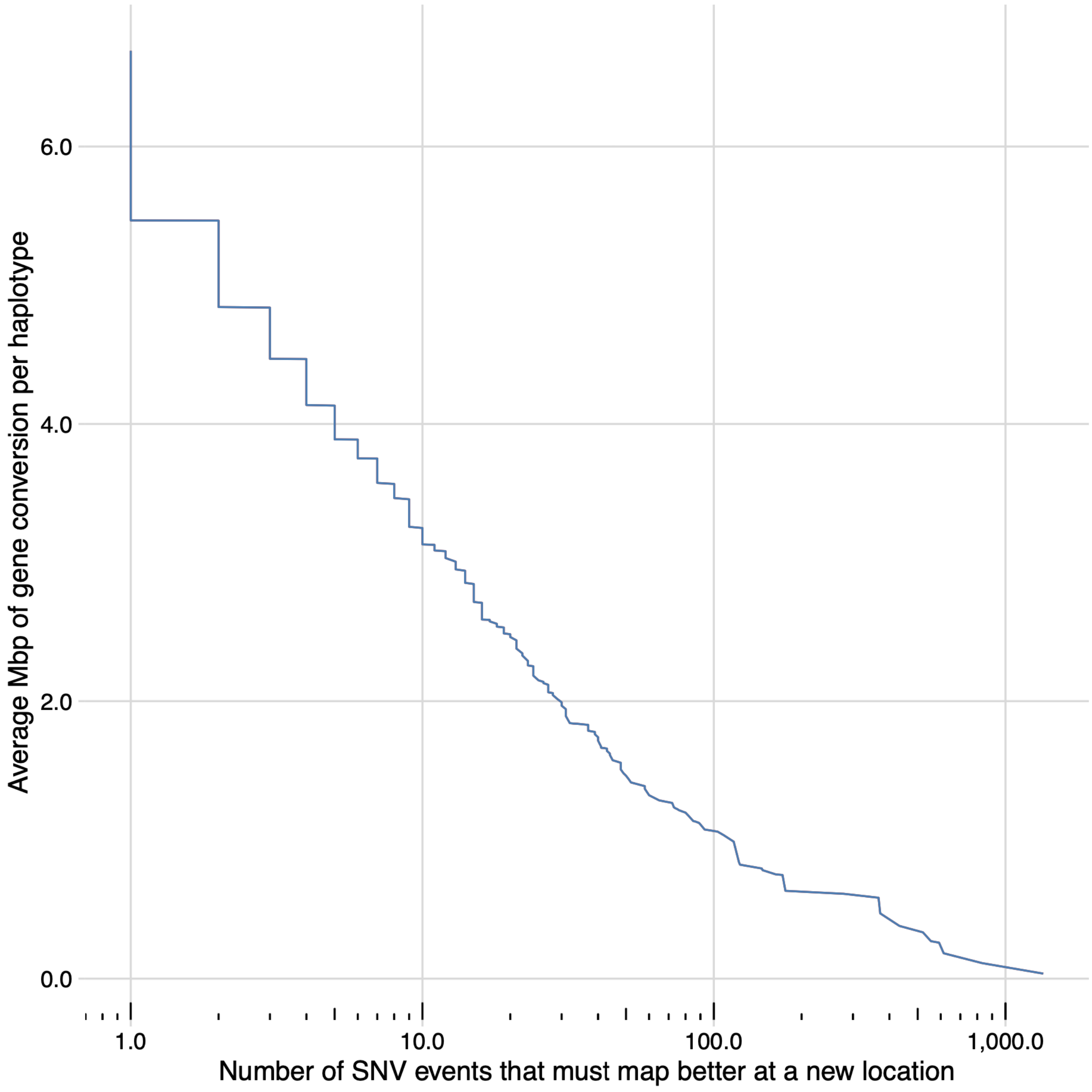


**Figure S3. Interlocus gene conversion in CHM1.** Mbp of IGC as a function of the number of supporting SNVs in CHM1.

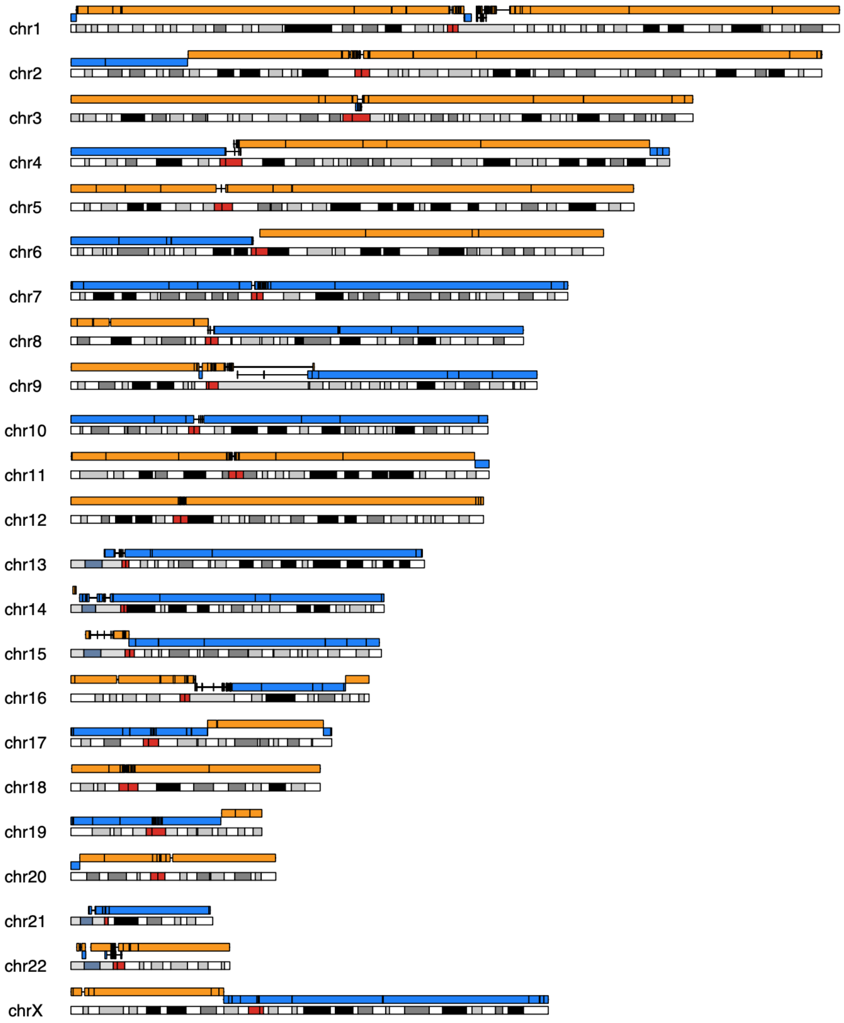


**Figure S4. Ideogram of an assembly of CHM1 aligned to T2T-CHM13.** The assembly of CHM1 has an N50 of 105.2 Mbp and allows for no accidental haplotype switching since it is from a hydatidiform mole, eliminating one significant mode of assembly error.

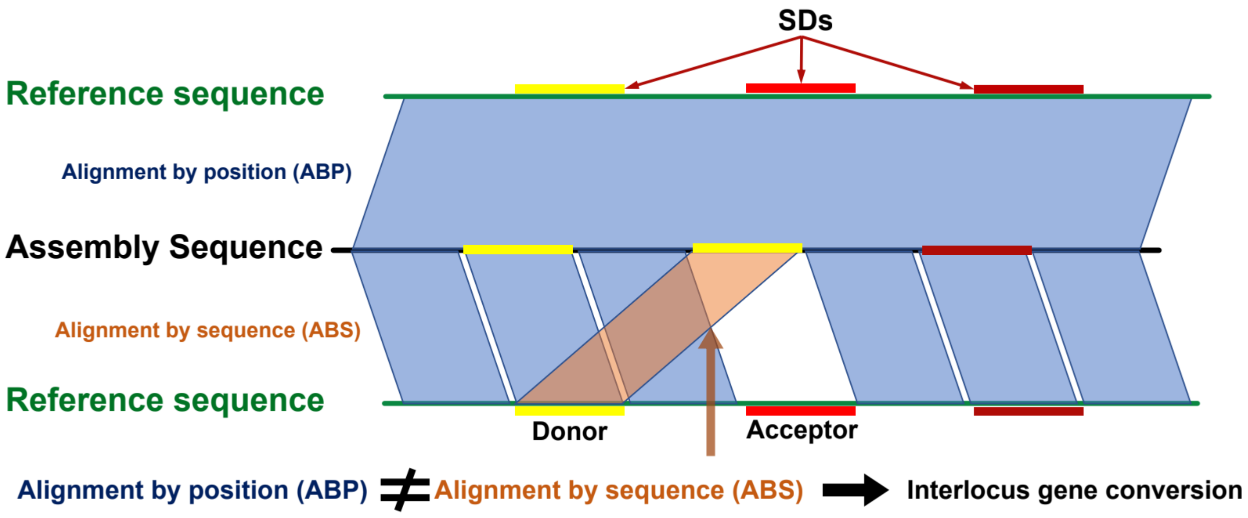


**Figure S5. Summary of methods for finding IGC.**

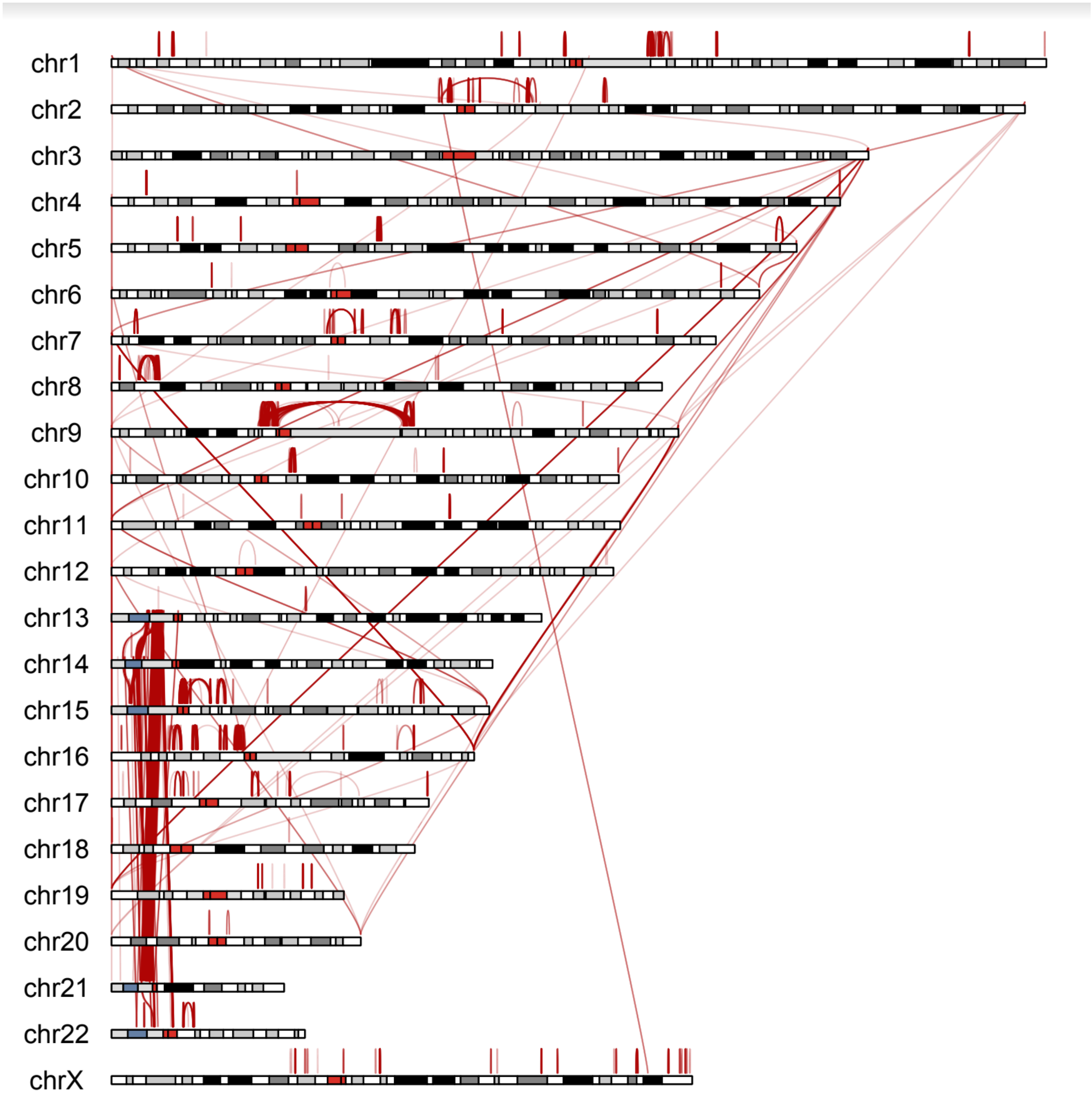


**Figure S6. Largest 10% of nonredundant IGC events in the genome.**

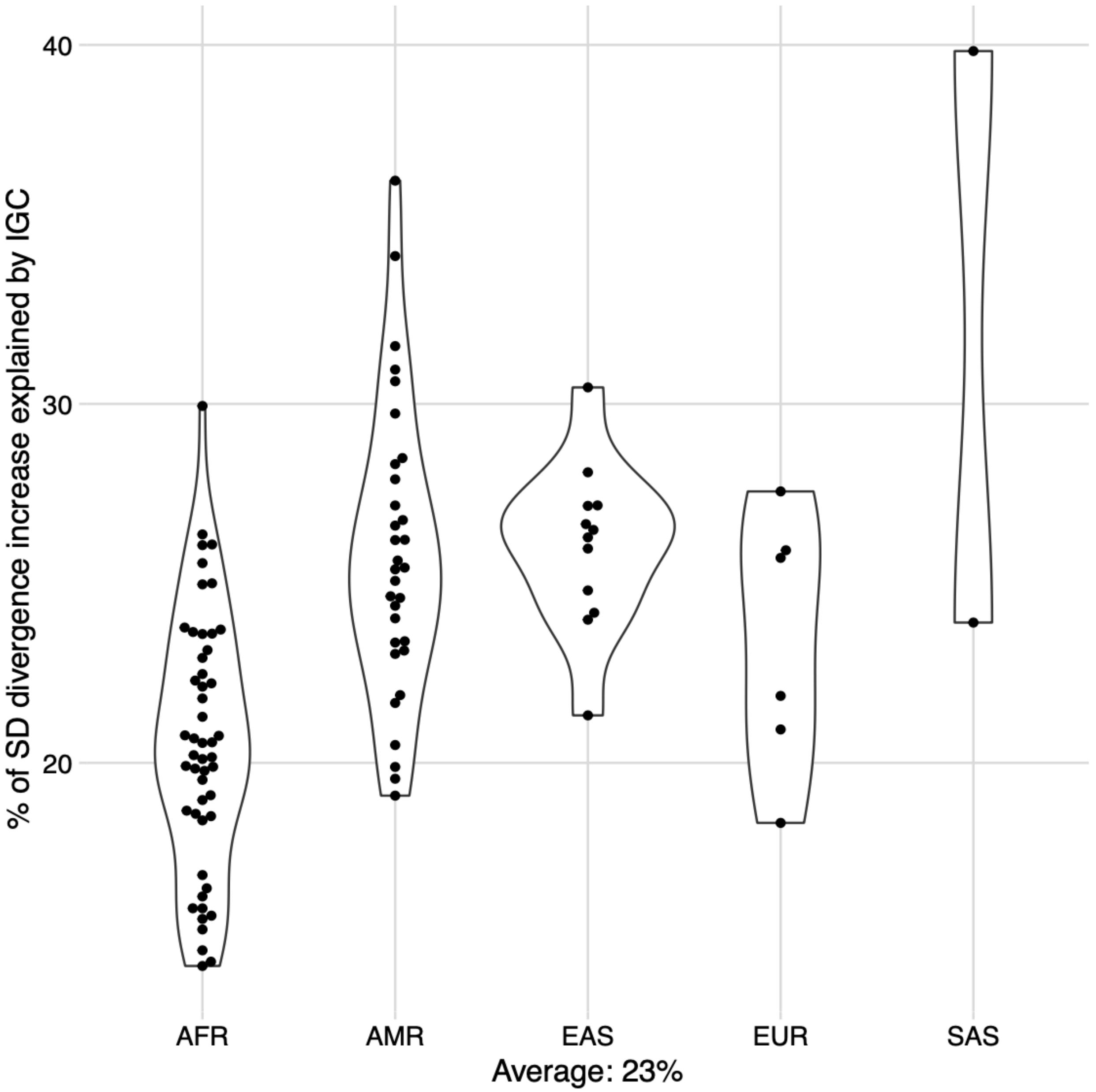


**Figure S7. Percent of increased single-nucleotide variation explained by IGC.** Here we calculate the average number of SNVs within SDs excluding regions of IGC and then take the difference from the original calculation to determine how much IGC contributed to the excess diversity in SNVs.

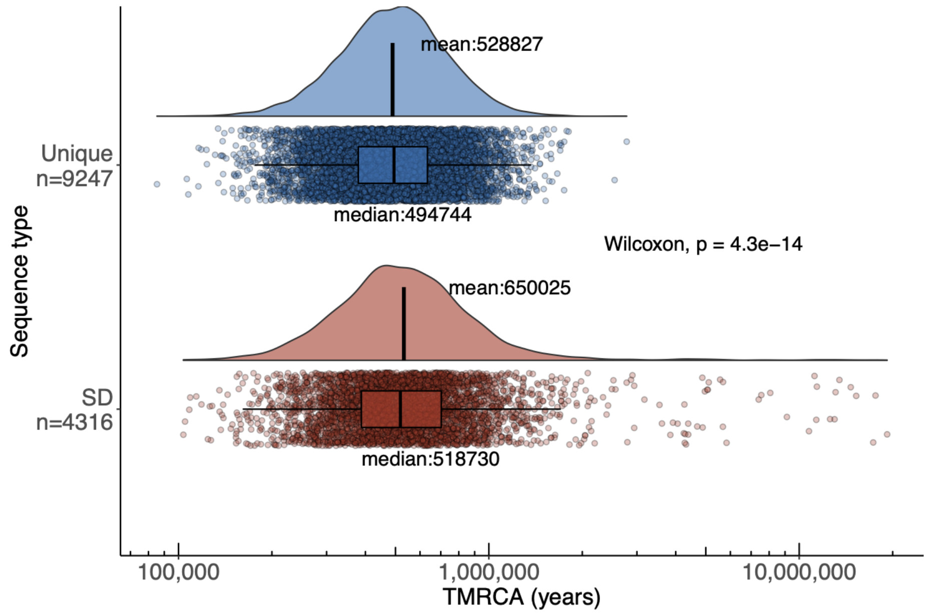


**Figure S8. Distributions of time to the most recent common ancestor (TMRCA) for unique (top) and SD (bottom) regions.** Measurements are based on nonoverlapping 10 kbp windows.

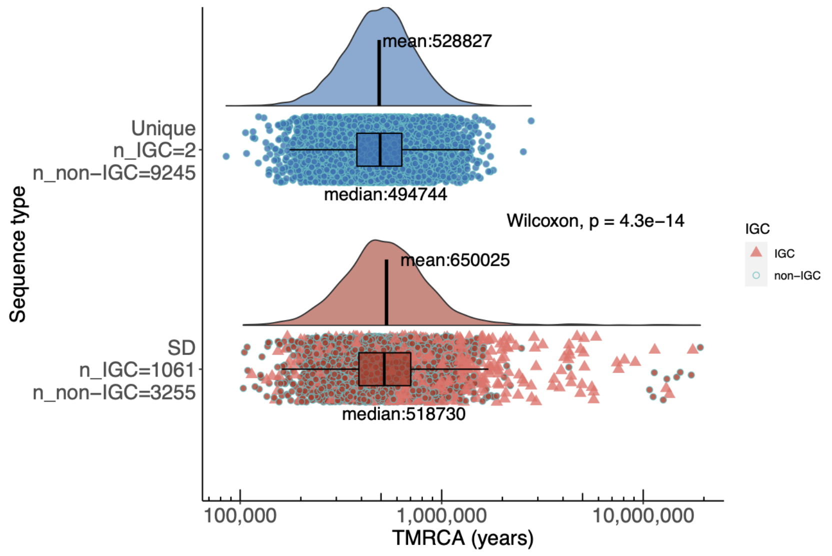


**Figure S9. Distributions of time to the most recent common ancestor (TMRCA) for unique (top) and SD (bottom) regions.** IGC sequences are marked as triangles.

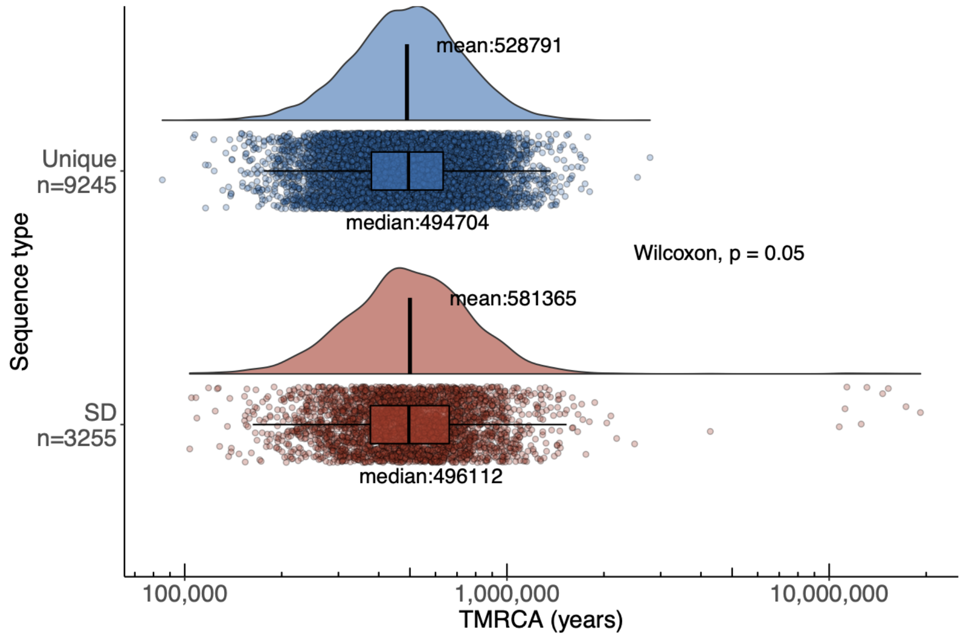


**Figure S10. Distributions of time to the most recent common ancestor (TMRCA) for unique (top) and SD (bottom) regions after excluding sequences affected by IGC.**

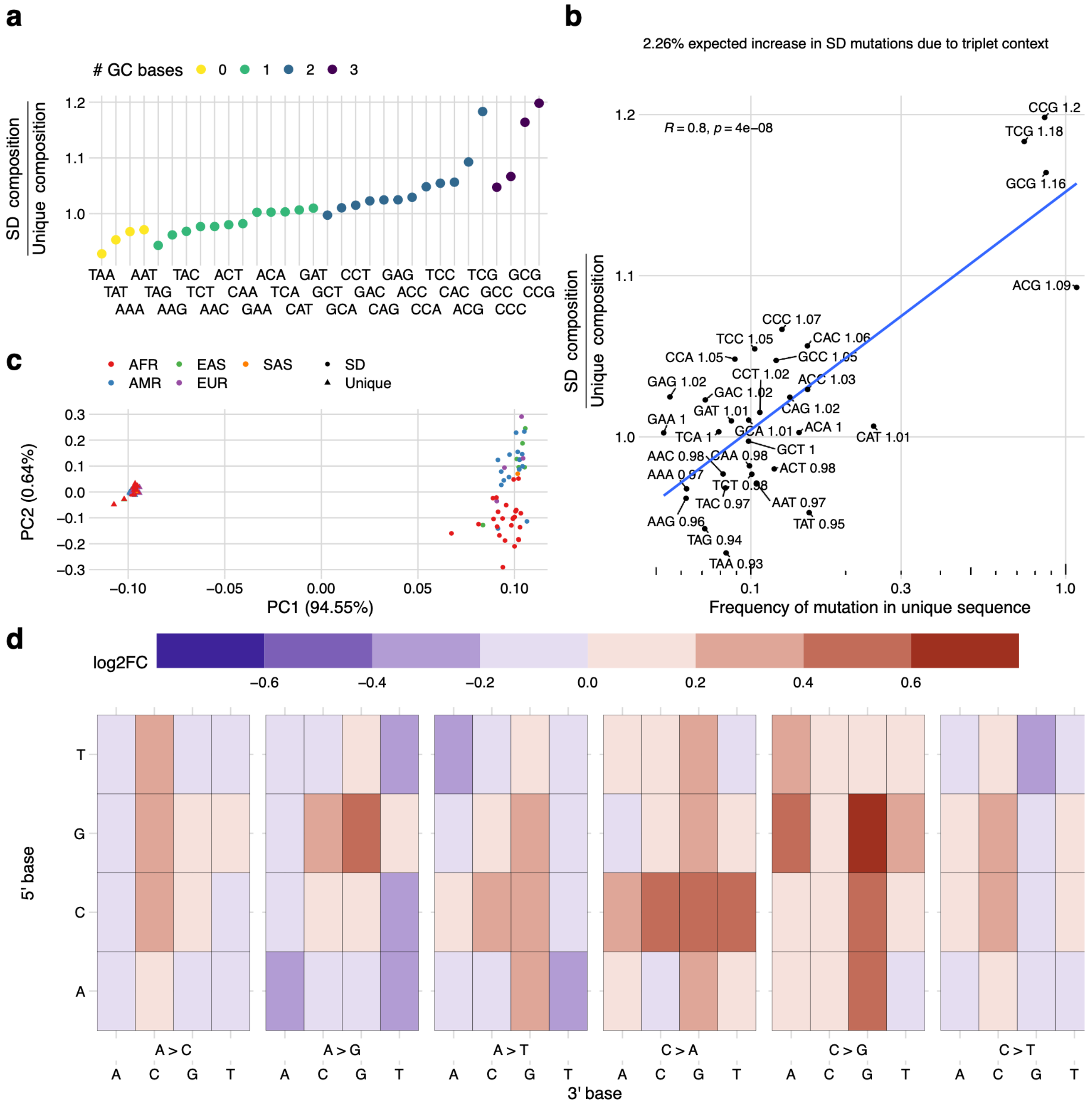


**Figure S11. Sequence composition and mutational spectra of SNVs in SDs without IGC.** **a)** Compositional increase in GC-containing triplets (3-mers) in SDs without IGC versus unique regions of the genome (colored by GC content). **b)** Shows a correlation between the enrichment of certain triplets in SDs compared to the mutability of that triplet in unique regions of the genome. Mutability is defined as the sum of all SNVs that change a triplet divided by the total count of that triplet in the genome. The enrichment ratio of SD over unique is indicated in text next to each triplet sequence. **c)** PCA of the mutational spectra of triplets in SD (circles) vs. unique (triangles) regions polarized against a chimpanzee genome assembly and colored by the continental superpopulation of the sample. **d)** Heatmap of the log-fold change in the triplet-normalized mutation frequency between SDs and unique sequences for the same mutational event.

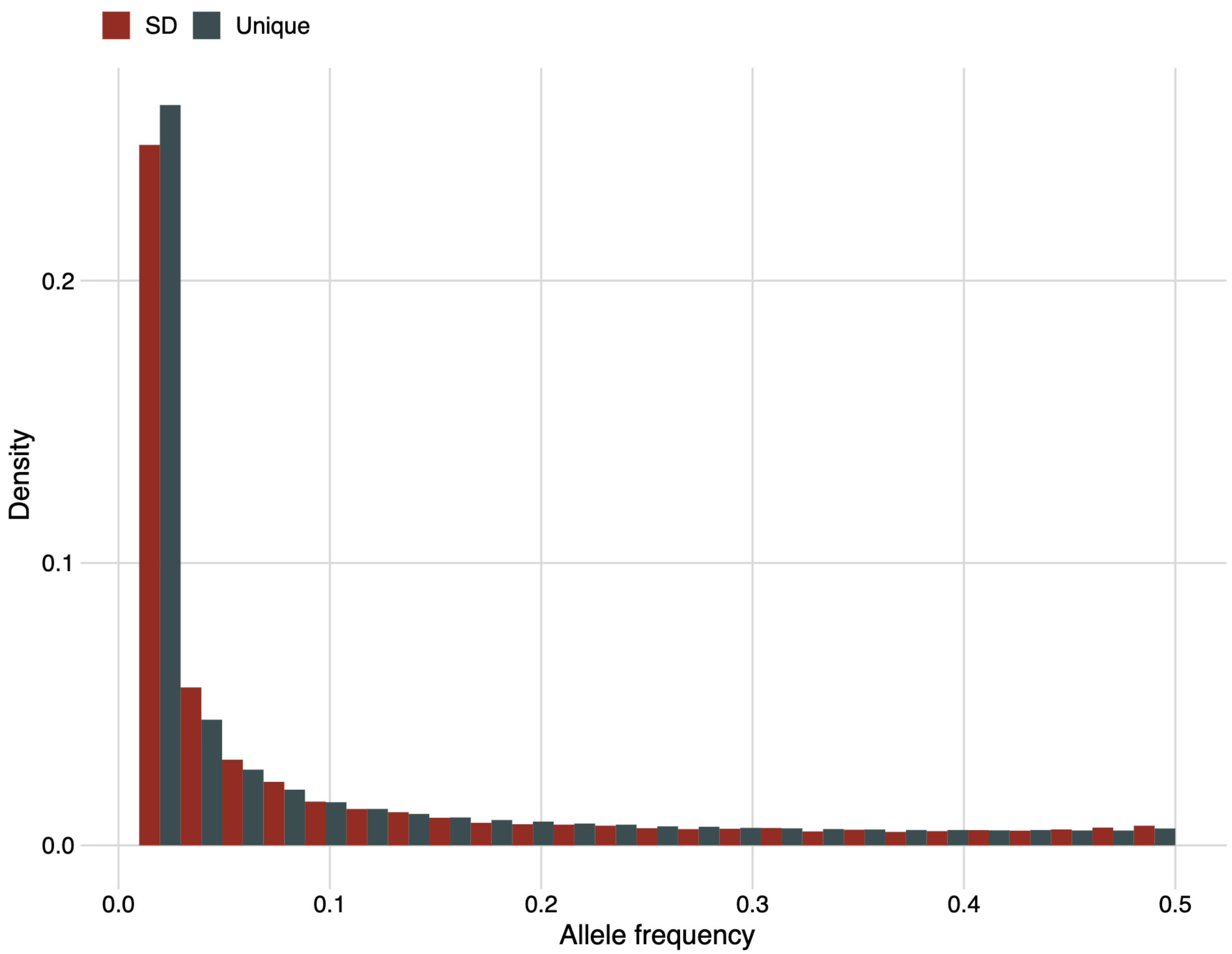


**Figure S12. Allele frequency spectrum for unique and SD SNVs.**

### Additional Figures

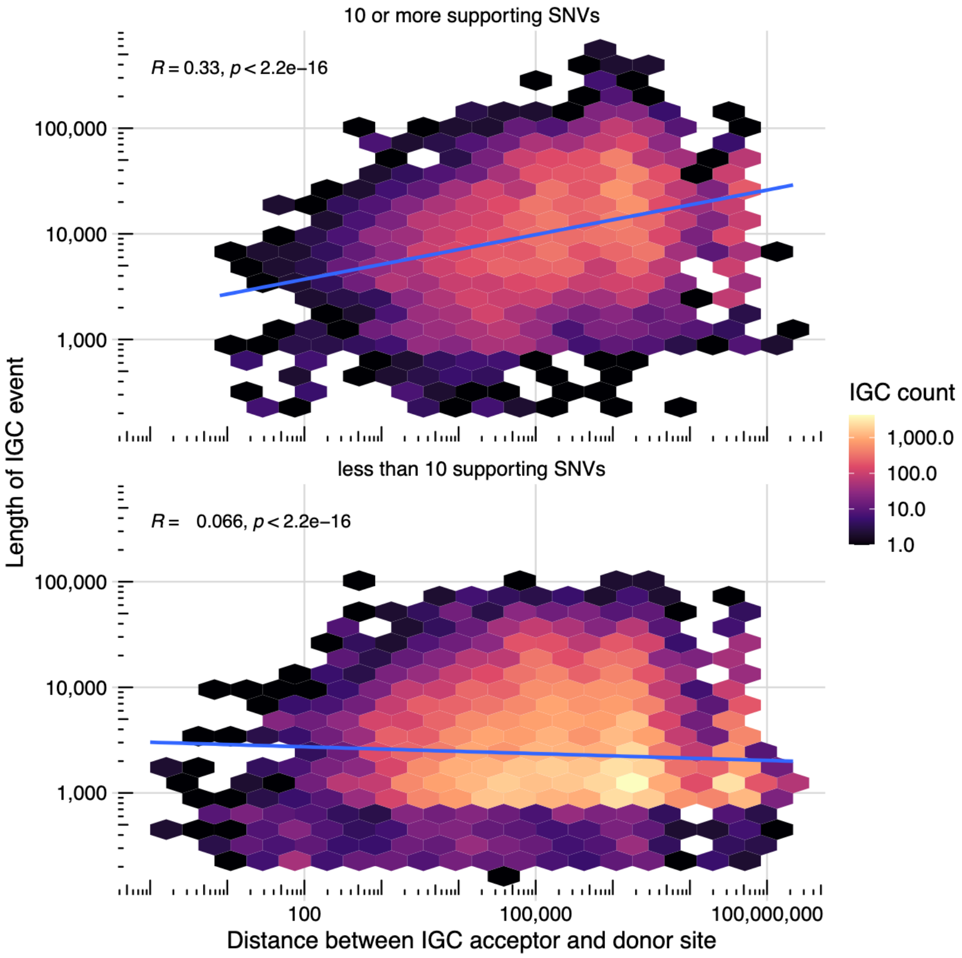


For events with strong evidence (10 or more supporting SNVs), we see a significant positive correlation (R=0.33, p<2.2e-16) between the length of the IGC event and the distance between donor and acceptor. This trend is not apparent in events with few supporting SNVs.

### References


Aitchison,J. (1982) The statistical analysis of compositional data. *J. R. Stat. Soc.*, **44**, 139–160.

Cheng,H. *et al.* (2021) Haplotype-resolved de novo assembly using phased assembly graphs with hifiasm. *Nat. Methods*, **18**, 170–175.

Dishuck,P.C. *et al.* (2022) GAVISUNK: Genome assembly validation via inter-SUNK distances in Oxford Nanopore reads. *bioRxiv*, 2022.06.17.496619.

Ebert,P. *et al.* (2021) Haplotype-resolved diverse human genomes and integrated analysis of structural variation. *Science*, **372**.

Koren,S. *et al.* (2018) De novo assembly of haplotype-resolved genomes with trio binning. *Nat. Biotechnol.*, **36**, 1174–1182.

Liao,W.-W. *et al.* (2022) A Draft Human Pangenome Reference. *bioRxiv*.

Logsdon,G.A. *et al.* (2021) The structure, function and evolution of a complete human chromosome 8. *Nature*, **593**, 101–107.

Rautiainen,M. *et al.* (2022) Verkko: telomere-to-telomere assembly of diploid chromosomes. *bioRxiv*, 2022.06.24.497523.

Sawyer GENECONV: a computer package for the statistical detection of gene conversion. *http://www. math. wustl. edu/~ sawyer*.
